# SHOT-R: A next generation algorithm for particle kinematics analysis

**DOI:** 10.1101/2021.12.21.473703

**Authors:** Eloina Corradi, Walter Boscheri, Marie-Laure Baudet

## Abstract

Analysis of live-imaging experiments is crucial to decipher a plethora of cellular mechanisms within physiological and pathological contexts. Kymograph, i.e. graphical representations of particle spatial position over time, and single particle tracking (SPT) are the currently available tools to extract information on particle transport and velocity. However, the spatiotemporal approximation applied in particle trajectory reconstruction with those methods intrinsically prevents an accurate analysis of particle kinematics and of instantaneous behaviours. Here, we present SHOT-R, a novel numerical method based on polynomial reconstruction of 4D (3D+time) particle trajectories. SHOT-R, contrary to other tools, computes *bona fide* instantaneous and directional velocity, and acceleration. Thanks to its high order continuous reconstruction it allows, for the first time, kinematics analysis of co-trafficked particles. Overall, SHOT-R is a novel, versatile, and physically reliable numerical method that achieves all-encompassing particle kinematics studies at unprecedented accuracy on any live-imaging experiment where the spatiotemporal coordinates can be retrieved.

## 1 Introduction

The recent advance of cutting-edge imaging techniques and the novel use of fluorescent labels over the last two decades have enabled to capture critical intracellular dynamic events in living samples. Along with these technical and experimental improvements, the advent of key analysis tools have been instrumental in studying these events, most notably to track moving particles’ direction and analyse velocity. However, the development of such algorithms has significantly lagged behind which prevents researchers from fully extracting information derived from live-imaging acquisitions (Fig. 1a) made by powerful instruments. For instance, it is currently not possible to compute important biological phenomena such as instantaneous velocity and acceleration of trafficked or co-trafficked cells or particles such as nanoparticles, viral particles, RNA and proteins (Fig. 1b). In addition, current approaches use approximations which yield poor accuracy to compute the velocity.

**Figure 1:**
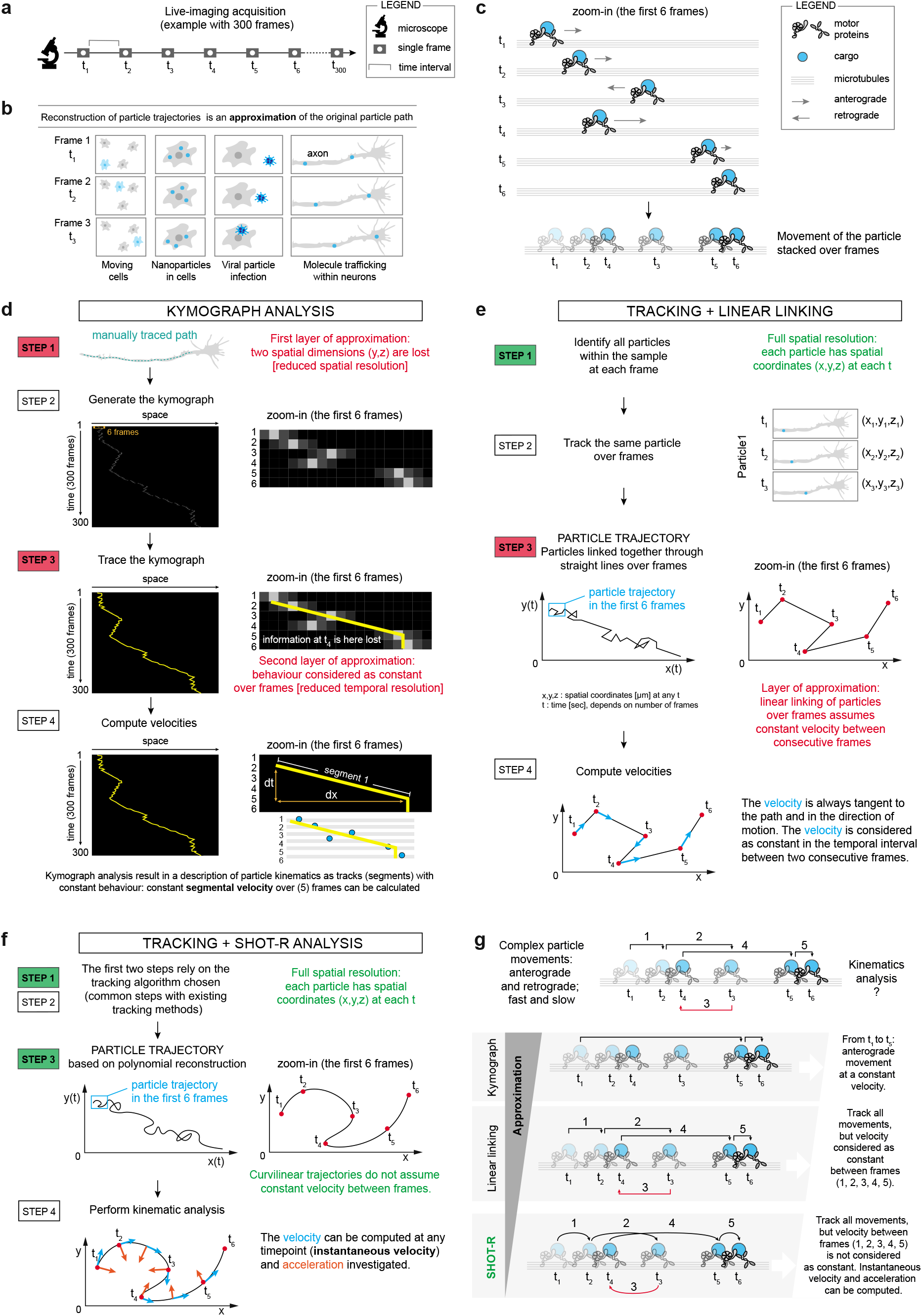
Particle kinematics analysis. **(a)** Schematic of a 300-frame live imaging experiment. **(b)** Examples of biological events studied by particle trajectory reconstruction, meaning by approximating the original particle path. **(c)** Example of the first acquired six frames (*t*_1_-*t*_6_) of a moving cargo transported by motor proteins on microtubules. Grey arrows indicate cargo directionality (e.g. from *t*_3_ to *t*_4_ the cargo moves retrogradely). **(d-f)** Schematic of the steps for particle kinematic analysis by kymograph (d) and tracking combined either with linear linking (e) or SHOT-R (f). Zoom-in panels matched the first 6 frames shown in (c). **(d)** Step 1 and 3 introduce approximations reducing both spatial and temporal resolution of the analysis. **(e)** In Step 1 the spatial resolution is preserved but in Step 3 the velocity is approximated, considering it as constant over consecutive frames. **(f)** SHOT-R by reconstructing the particle trajectory with polynomials, does not assume constant velocity between frames, increasing the spatiotemporal resolution and allowing to investigate velocity and acceleration at any point of the trajectory. **(g)** SHOT-R drastically reduces the approximation of the particle trajectory reconstruction, improving the overall accuracy in the analysis. Abbreviations: SHOT-R, Spatiotemporal High Order Trajectory Reconstruction.

Kymographs are largely used to investigate particle velocity [1, 2, 3, 4, 5]. Kymographs are 2D graphical representations of the spatial position of an object (particle etc.) over time in which the *x* axis represents one spatial dimension (instead of three) and the *y* axis represents time. Kymographs are obtained by stacking serial frames of an acquisition (Fig. 1a,b). In this stack, only a small portion of each frame is selected by tracing a manually defined path, in effect losing information pertaining to *y* and *z* spatial dimensions. Particle velocity is computed by tracing a line over the trajectory of an object when it appears grossly constant over multiple serial acquisitions (Fig. 1c,d). The slope of the traced line represents the particle velocity along that given path (*dx*_*s*1_/*dt*_*s*1_) (Fig. 1d; Extended Data Fig. S1a). Overall, kymograph analysis oversimplifies the complexity of a tracking problem by assuming that the trajectory of a particle is constant over several acquisitions and by selecting a one dimensional fixed spatial path. As a consequence, both spatial and temporal resolutions are strongly reduced (Fig. 1c,d; Extended Data Fig. S1a).

In addition to kymographs, single-particle tracking (SPT) methods have also been employed in the field to analyse cell and particle velocity (Fig. 1c,e). SPT is a more advanced and accurate approach than kymographs and is based on a two-step process: i) detection of particle spatial position in three dimensions (*x, y, z*) frame by frame and ii) linking particle spatial position with straight lines over time between consecutive frames to define the particle trajectory [6, 7] (Fig. 1c,e). The critical challenge of this approach is to be able to identify and track the same particle from one frame to the next. To this end, several approaches have been developed over the years depending on the type of obtained acquisition. Particle detector and linking algorithms can be chosen according to the difficulty that the acquisition poses (e.g. high particle density, low signal-to-noise ratio, motion heterogeneity, blinking signal, particle merging or splitting) [8, 9, 10, 11].

Contrary to kymographs, SPT allows full spatial resolution because *x, y* and *z* coordinates are computed at each time point t. The temporal resolution is, however, reduced. Due to the time-lapse nature of an acquisition, no information about particle position is known between frames. The linking process assumes that the particle velocity is constant between consecutive frames whilst it may not be. Under this assumption, a mean velocity is calculated between frames (or ”frame to frame velocity”) but instantaneous velocity or acceleration cannot thus be computed (Extended Data Fig. S1a,b). While SPT provides significant improvements in both spatial and temporal resolution over kymographs, its inherent assumption of constant particle velocity constitutes a critical limiting step in image analysis because it overlooks particles showing rapid changes in velocity, a commonly observed phenomena in biology, and precludes the accurate analysis of particles’ co-trafficking. While massive efforts have been invested for improving the particle detection and linking of SPT [12, 13, 7, 14, 15, 16, 17], an entirely new approach that tackles and overcomes the basic erroneous assumption of constant velocity is direly needed to better match the true nature and complexity of particle kinematics.

Here, we have developed a next generation algorithm, SHOT-R (Spatiotemporal High-Order Trajectory Reconstruction), that goes beyond SPT algorithm performance and accuracy. Through SHOT-R, particles’ spatial positions are tracked and linked with currently available SPT algorithms but instead of linking them with segments (”linear linking”), SHOT-R links them with curvilinear trajectories thanks to a polynomial reconstruction (Fig. 1c,f,g). This approach does not assume constant velocity between frames and as a consequence, finally allows to compute particle instantaneous velocity and acceleration at any time points, including between frames, and to study the kinematics of the co-trafficking of multiple particles at once (Extended Data Fig. S1a,b). Overall, this novel approach preserves the full spatial resolution of particle tracking and significantly improves the temporal resolution over SPT (Extended Data Fig. S1c,d).

## 2 Results

### 2.1 SHOT-R reconstructs trajectory through polynomials

Particle kinematics is described by ordinary differential equations (ODE) governing the motion:

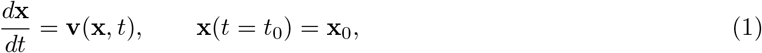

where *t* denotes the time coordinate, **x** = (*x, y, z*) is the position vector of spatial coordinates and **v**(**x**,*t*) = (*v_x_, v_y_, v_z_*) is the velocity vector with components along each direction in space. The trajectory starts at the initial time *t* = to from the position **x**_0_ (Fig. 2a) and the ODE Eq.(1) describes the motion of the particle over time. The trajectory equation Eq.(1) is a continuous function, whereas experimental data only provide a discrete set of known spatial coordinates **x**_*k*_ at the corresponding time *t_k_*, since imaging is performed through time-lapse acquisitions and is thus inherently not continuous (Fig. 1a). The reconstruction of particle trajectories, therefore, merely represents an approximation of the original particle trajectory.

**Figure 2:**
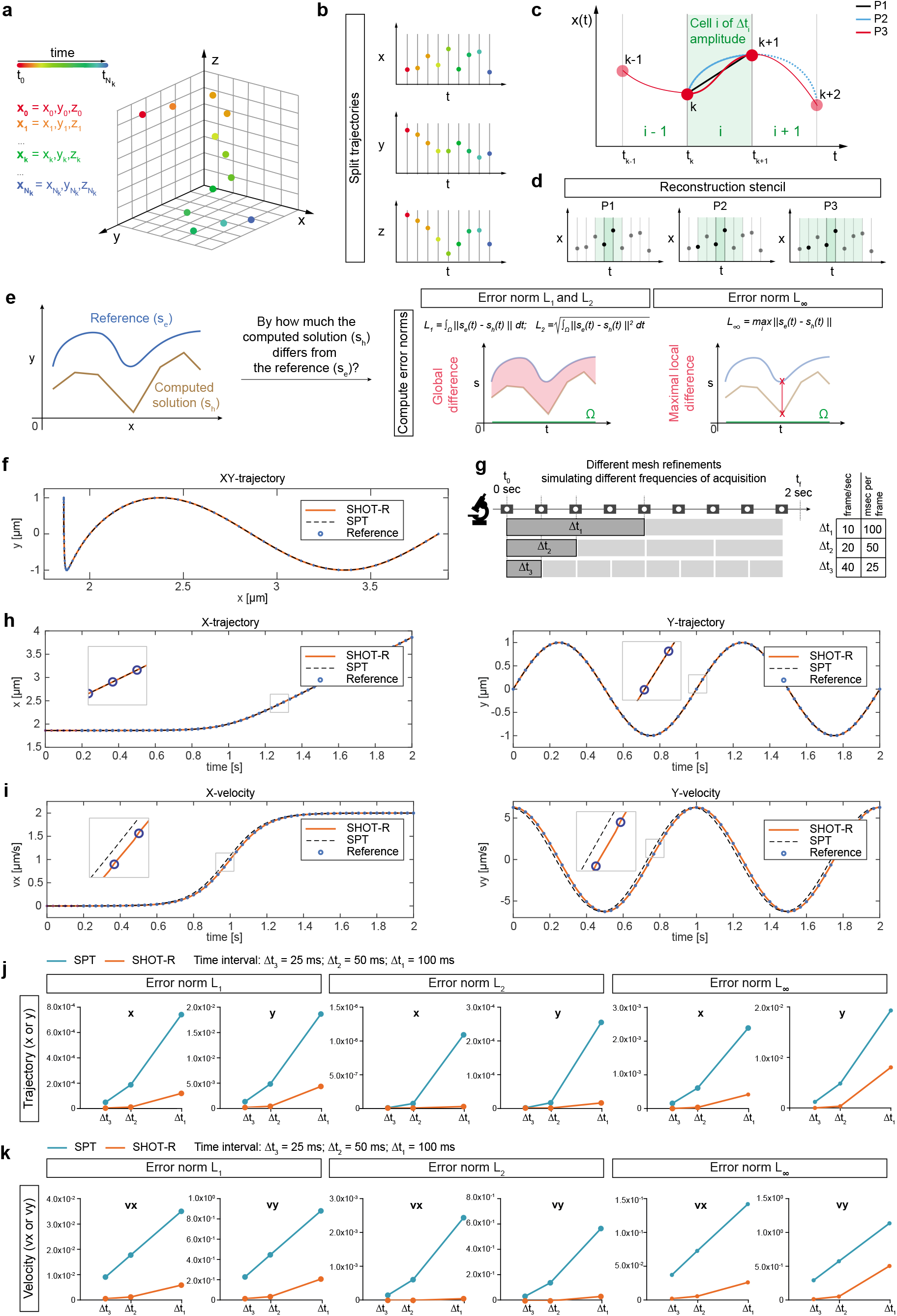
Spatiotemporal high order trajectory reconstruction (SHOT-R) method and its mathematical validation. **(a)** Schematic of SHOT-R 4D input data: particle motion is described as the evolution of the spatial coordinates (*x, y, z*) in time (from *t*_0_ to *t_N_k__*). **(b)** Schematic of space-time split trajectory strategy. Each direction (*x, y, z*) is analysed separately, thus a multidimensional setting can be reduced to multiple one-dimensional problems. **(c,d)** Schematic of trajectory reconstruction based on polynomials (SHOT-R). An interpolation polynomial *p^N^*(t) of arbitrary degree *N* is constructed for each cell *i* (c) by considering the cell itself and a total number *N* + 1 surrounding elements (i.e. the reconstruction stencil) (d). **(e)** Schematic representation of error norms calculation. Eq.(S39). **(f,g)** Parameters used in the new mathematical test: XY-reference trajectory mimicking particle movement (f) and different mesh refinements simulating timelapse experiments at different acquisition frequency (g). **(h-k)** Qualitative (h,i) and quantitative (j,k) comparison between SPT and SHOT-R in trajectory reconstruction (h,j) and in computing velocity (i,k) on given equations Eq.(S37)-(S38). (f,h,i) Δ*t*_2_ mesh refinement. Abbreviations: *P*1,*P*2, *P*3, first, second, third degree polynomial; Δ*t*, time intervals (mesh refinement); SPT, single particle tracking.

Current SPT methods approximate the original trajectory by a first degree polynomial reconstruction (i.e. linear linking, Fig. 1e), while SHOT-R algorithm solves particle kinematics equations Eq.(1) based on higher order polynomials (Fig. 1f). Contrary to SPT, SHOT-R reconstructs trajectory with a greater accuracy and therefore better reflects the inherent particle motion. SHOT-R input data are the spatiotemporal coordinates of particles retrieved from the raw movies. These particles move over time in a 3D space and therefore have 4 dimensions at any given point (*x, y, z, t*). Reconstructing the particle 3D trajectory is computationally demanding because it requires a complex 3D spatial mesh at each timepoint. We tackle this challenge by splitting the 3D trajectory into multiple one-dimensions (Fig. 2b), thereby solving in parallel three [*x*(*t*), *y*(*t*), *z*(*t*)] one-dimensional equations Eq.(1), which are then combined to obtain the position of the particle at any time point given by a 4D vector in (*x, y, z, t*). Overall, this approach speeds up and simplifies the computational process without losing information on the actual particle position.

In parallel with the step described above, each 1D trajectory is reconstructed with a high order polynomial. First, the computational mesh is built as follows. For each dimension, each cell *i* is limited in time by two points of the trajectory (*t_k_*, *t*_*k*+1_) and has length Δ*t_i_* (Fig. 2c). Second, the 1D trajectory is reconstructed step-wise locally within a given cell using information of the particle position from adjacent cells (Fig. 2c). The assembly of these contiguous cells forms a stencil whose size depends on the polynomial degree used for the reconstruction (Fig. 2d). Specifically, the stencil size comprises a total number of cells which is at least equal to the number of degrees of freedom of the chosen polynomial degree. Therefore, the higher the polynomial degree the wider the stencil, i.e. the more information from neighbouring cells is required for the reconstruction. SHOT-R thus requires more information (more datapoints) than SPT to be able to reconstruct trajectories. Third, a constraint is applied to ensure that the reconstructed trajectory passes through the actual particle positions by imposing the exact interpolation of the input coordinates **x**_*k*_.

In practice, SHOT-R final result is a piecewise continuous polynomial function of degree *N* which reconstructs at order 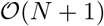 the trajectory along each spatial dimension (*x, y, z*).

### 2.2 Mathematical tests validate the accuracy and the robustness of SHOT-R

We next validated the degree of accuracy of SHOT-R through a mathematical-driven approach. First, we performed a convergence study [18] (for details refer to Method section S1.7). Such convergence analysis reveals whether the numerical solution provided by a numerical method (here, SHOT-R) approaches the exact solution of a standardized analytical equation Eq.(S36). It does so by computing the difference (error norms, Eq.(S39), Fig. 2e) between the two solutions at specific time intervals Δ*t* (mesh refinement). Such a difference is expected to decrease at a given rate according to the polynomial degree and the temporal interval applied: the higher the polynomial degree and the smaller the temporal interval, the lower the measured difference with the exact solution and thus the higher the accuracy of the numerical method. We demonstrated up to the fifth order that SHOT-R fulfilled this requirement, proving its numerical accuracy (Supplementary Table S1).

Second, we developed a new mathematical test to assess SHOT-R precision and improvement specifically over current SPT approaches by using equations mimicking particle movement.

To achieve this, we first constructed through known functions (see Eq.(S37)-(S38) in Methods), an hypothetical reference xy-trajectory of a particle with a complex behaviour: along the *x*-axis the particle stalls, then accelerates and eventually moves at a constant velocity while along *y*-axis it oscillates regularly (Fig. 2f). Three different mesh refinements were considered (Δ*t*_1_, Δ*t*_2_ and Δ*t*_3_), simulating increasing frequencies of acquisition (increased input points *x_k_*) and thus increasing temporal resolution (Fig. 2g). Next, the particle trajectory was reconstructed and the velocity computed with both SPT and SHOT-R (Fig. 1e,f). For the latter, we chose to use a third degree polynomial as a compromise between accuracy and efficiency, since the higher the polynomial degree, the higher the accuracy (see above) but the higher the computational cost. While with an intermediate mesh density (Δ*t*_2_) the trajectory was overall appropriately reconstructed by both SPT and SHOT-R compared to the reference (Fig. 2h), the velocity vector was more accurate with SHOT-R (Fig. 2i). Importantly, *v_x_* computed through SPT was not accurate specifically where the velocity changed (Fig. 2h, boxed).

Furthermore, we measured *L*_1_, *L*_2_ and *L*_∞_ norms (Eq.(S39), Fig. 2e) and SHOT-R systematically achieved better results than SPT in estimating particle trajectory and velocity, with errors of about two orders of magnitude smaller with respect to SPT (Supplementary Table S2, Fig. 2j,k). This effect was dependent on the simulated acquisition frequency itself, and as expected the shorter the time interval the smaller the errors for both SHOT-R and SPT. Nevertheless, already at the intermediate frequency of acquisition (20 frame/s; Δ*t*_2_ in Fig. 2f), the error in computing *v_y_* with SPT method was 0.44 *μm/s* contrary to the 0.02 *μm/s* using SHOT-R. In conclusion, the mathematical tests validate the accuracy and the robustness of SHOT-R, and show its superior capabilities compared to SPT in computing velocity, thus making it, on principle, suitable and reliable for applications on real biological datasets.

### 2.3 SHOT-R increases accuracy in computing velocity on biological datasets

We next investigated whether SHOT-R performs better than SPT on a biological dataset we recently published, namely endogenous miRNA (pre-miR-181a-1) trafficking along axon [5] (Extended Data Fig. S2a). Previously, pre-miR-181a-1 kinematics was studied by kymograph analysis [5], while here, we retrieved particle coordinates with TrackMate [19] (Extended Data Fig. S2a-c) and applied SHOT-R or SPT. SHOT-R and SPT trajectory reconstruction approaches were compared. As already observed for the mathematical test (Supplementary Table S2, Fig. 2j), biological particle trajectories did not change depending on the degree of the polynomial reconstruction (SPT: first degree vs SHOT-R: third degree polynomial) (Fig. 3a-c, Extended Data Fig. S2d). Indeed, the mean values of norm distribution (Fig. 2e, for definition) were close to zero (i.e. *L*_1_, 0.08 *μm*; *L*_2_ and *L*_∞_, 0.03 *μm*), suggesting a perfect overlap of the SPT and SHOT-R reconstructed trajectories (Fig. 3c). On the contrary, the computed velocities differed (Fig. 3d-f) with mean errors greater than zero (*L*_1_, 0.65 *μm/s*; *L*_2_, 0.24 *μm/s*; *L*_∞_, 0.26 *μm/s*) (Fig. 3f).

**Figure 3:**
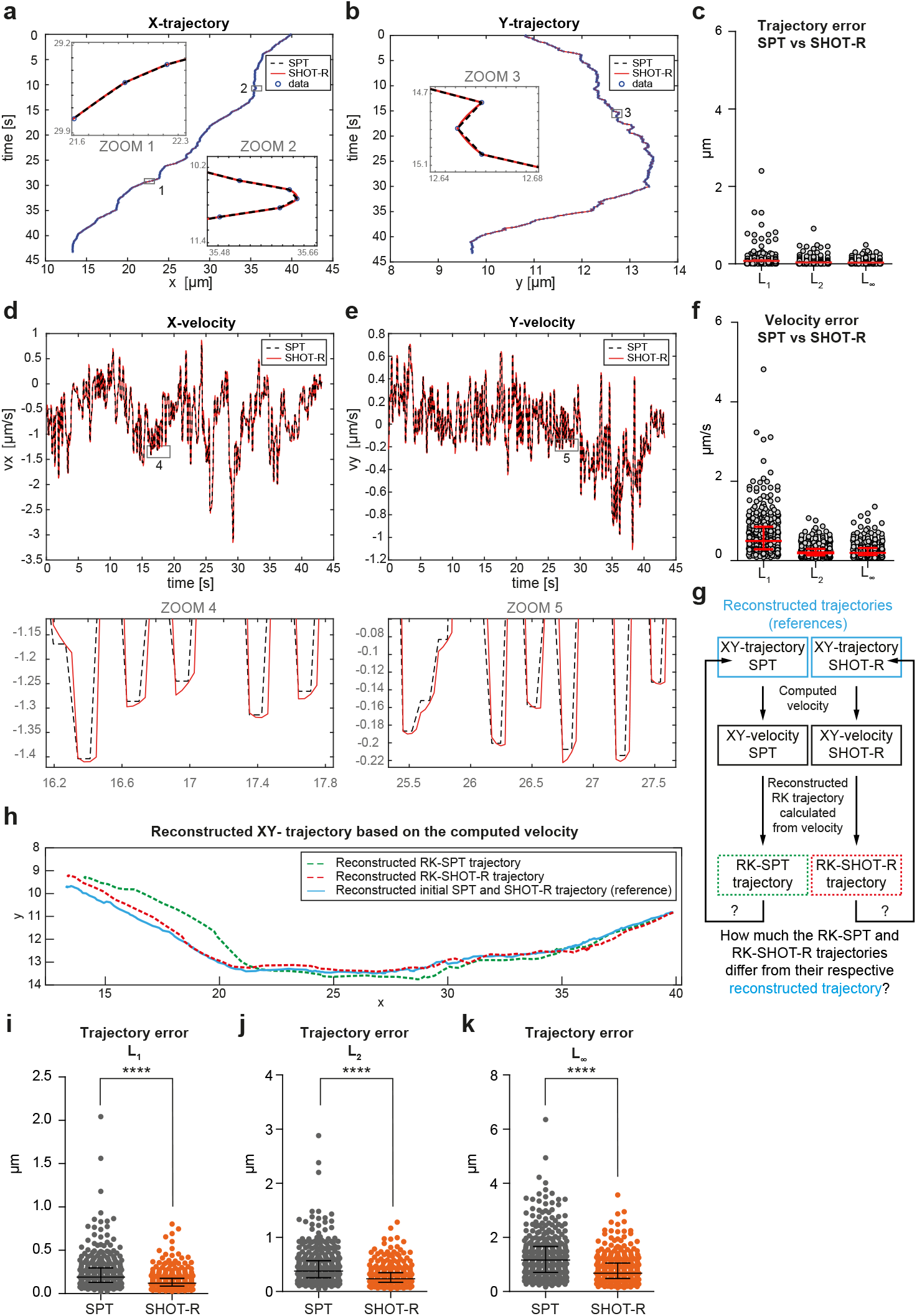
SHOT-R increases accuracy in computing velocity on a biological dataset (a,b,d,e) Representative comparison of SPT and SHOT-R in *xy* trajectory reconstruction (a,b) and in computing velocity (d,e). **(c,f)** Difference between SPT and SHOT-R reconstructed trajectory (c) or computed velocity (f) measured as error norms (*L*_1_, *L*_2_, *L*_∞_). **(g)** Schematic of the pipeline to test SPT and SHOT-R accuracy in calculating velocity. Color code matches panel h. **(h)** Representative comparison of RK-SPT and RK-SHOT-R trajectory obtained by backward integration of the reconstructed velocity. **(i-k)** Difference between RK-SPT and RK-SHOT-R measured as error norms *L*_1_ (i), *L*_2_ (j), *L*_∞_ (k) of the reconstructed trajectory based on velocity trace. Abbreviations: data, the **x**_*k*_ = (*x_k_,y_k_*) coordinates retrieved with TrackMate; RK, Runge-Kutta method for backward time integration. Data information: **** P < 0.0001. Values are median with interquartile range. Data are not normally distributed (Shapiro-Wilk test), two-tailed Wilcoxon matched pairs test. Each data point corresponds to a particle. Total number of analysed particles: 372 (c, f, i-k).

These errors are remarkable considering that the velocity of moving axonal particles range from 0.2 *μm/s* to 0.5 *μm/s* (slow-movements) and more than 0.5 *μm/s* (fast movements) [5, 20, 21]. As a consequence, the error produced by the current available method could alter the biological interpretation of the data with the risk of drawing wrong conclusions (i.e. over- or under-estimation of the real particle kinematics).

While this approach enabled us to compare SHOT-R to SPT, it didn’t allow us to analyse which of the two methods better approximates the real biological dataset, given the lack of an exact reference for the velocity. We thus implemented a novel test to assess whether SHOT-R is significantly more accurate in computing velocities than available strategies by using an internal reference to which reconstructed trajectories are compared (Fig. 3g). Through this test, we aimed to evaluate the errors in computing velocity between each method (SHOT-R or SPT) their respective internal reference. The smaller the error, the most accurate the method. To do this, *xy* trajectories of all particles were firstly reconstructed with first and third polynomial degree based on **x**_k_ coordinates of all tracked particles. Second, velocity in xy was computed from SPT and SHOT-R reconstructed trajectories. Third, the particle trajectories ”RK trajectories” generated from the velocity were built by backward time integration of the ODE Eq.(1) through Runge-Kutta (RK) methods (see method section S1.10 for details). In other words, while velocity is usually computed from the trajectory, we did the opposite in this step: we reconstructed the trajectory starting from the calculated velocity. Since the velocity is the derivative of the position with respect to time, the RK-trajectories were obtained by integrating in time the velocity (i.e. backward time integration). Lastly, the RK-SPT and RK-SHOT-R trajectories were compared to their respective reconstructed trajectory (the references) given by the known data points **x**_*k*_ (Fig. 3g,h). We found that the error generated by SHOT-R was significantly less compared to the one generated with SPT approach (Fig. 3i-k), both considering the curve globally (*L*_1_ and *L*_2_, Fig. 3i,j), as well as the maximal difference in position obtained for each trajectory (*L*_∞_, Fig. 3k). This data suggest that SHOT-R is more precise than SPT in estimating particle velocities. In summary, similarly to the mathematical test, SHOT-R drastically increases the accuracy in computing velocity on a real biological dataset.

### 2.4 SHOT-R provides novel insight on particle kinematics: from average to instantaneous velocity and acceleration

Having validated the theoretical performance of our new method, we next explored whether SHOT-R enables us to gain greater biological insight on particle kinematics (i.e. average and instantaneous bahaviour) exploiting our pre-miRNA axonal trafficking dataset [5]. To obtain an overview on the particle motility, we first investigated the average speed (*v_L_*, Eq.(S26)) and the average velocity both based on global (*v_D_*, Eq.(S27)) or local (*v_M_*, Eq.(S28)) displacement (Extended Data Fig. S1a and S3a). The average speed depends on the actual distance traveled by the particle (Extended Data Fig. S3a). Trajectory reconstruction by SPT and SHOT-R differ, as they are based on first versus higher order polynomial. For SPT, this distance is by definition the shortest path possible whilst for SHOT-R, it is close to the actual distance traveled by the particle as demonstrated above in Section 2.2. Unsurprisingly, we found that the average speed of axonal pre-miRNA computed with SHOT-R and with the SPT significantly differed in a paired comparison test (Extended Data Fig. S3b, mean values: 0.6943 *μm/s* vs 0.6971 *μm/s*). However, although significant, such a small difference does not change the biological conclusion that might be drawn for the analysed dataset. The average velocity instead depends on particle positions at given time of acquisition. Both SPT and SHOT-R reconstruct particle trajectories from positions determined by an identical tracking algorithm so, again, as expected, axonal pre-miRNA average velocities were identical for both methods (Extended Data Fig. S3c,d). Overall, this shows that SHOT-R performs better than SPT to compute speed but not average velocities.

We, then, deeper investigated particle kinematics at higher temporal resolution, by computing particle (axonal pre-miRNA) mean velocity between consecutive frames i.e. ”frame to frame velocity” along each trajectory (Extended Data Fig. S1a). Such analysis is only possible with SPT and SHOT-R but not with kymograph (Fig. 1d). The particle velocity calculated for each portion of trajectory between two frames (Extended Data Fig. S1a) was classified as stationary, slow and fast moving as previously published [5, 20, 21] (Fig. 4a,b). We found that particles moved predominantly with fast frame to frame velocities along trajectories (46.6% fast moving compared to 35.1% slow moving and 18.3% stationary) (Fig. 4b). This approach also enabled us to determine the proportion of the trajectory for which particles were in movement. We detected that particles moved at least along a third of the trajectory whilst remaining immobile for the remaining two third. Some particles even exhibited fast movement behaviour along the entire length of the trajectory. Overall, this suggests that pre-miRNAs are highly motile molecules. Such a result is in line with our previous analysis showing that pre-miRNAs were preferentially actively transported along axon (Mean Square Displacement analysis, which relies on SPT) at an average velocity based on total displacement of 0.24 *μm/s* (kymograph analysis) [5]. However, here, the frame to frame velocity analysis revealed that even particles with low average velocity present some fast movements (not a single pre-miRNAs puncta showed stationary behaviour along the entire trajectory, Fig. 4b). This demonstrates that particles have a more complex behaviour than our previous analyses were able to reveal.

**Figure 4:**
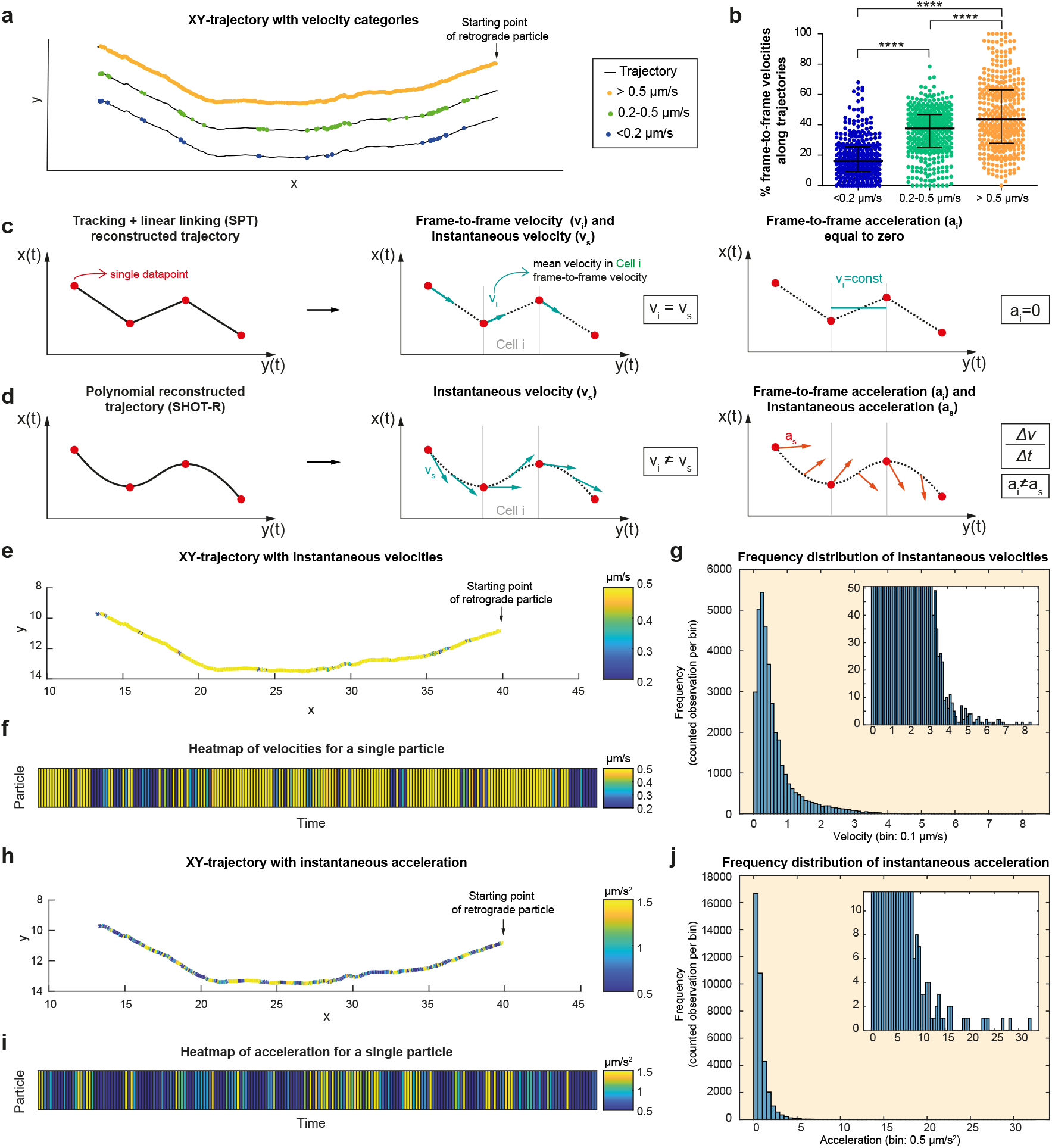
SHOT-R: from average to instantaneous velocity and acceleration. **(a)** Representative xy trajectory of a single moving particle (same as panel (e) and (h)). Single colored dots on the trajectory represent the category of frame to frame velocities *v_i_*. **(b)** Percentage of frame to frame velocity category along each particle trajectory. **(c,d)** Schematics of velocity and acceleration retrieved from linear (c) or polynomial (d) reconstruction of the trajectory. **(e,h)** Representative xy trajectory. Colors represent the magnitude of instantaneous velocities (color bar from 0.2 *μm/s* to 0.5 *μm/s)* (e) or the magnitude of instantaneous acceleration (color bar from 0.5 μm/s^2^ to 1.5 μm/s^2^) (h). **(f,i)** Heatmap of the instantaneous velocities (f) or acceleration (i) over time for the particle in panel (e) and (h). **(g,j)** Frequency distribution of instantaneous velocities *v_s_* (g) or acceleration *a_s_* (j) of all analysed particles. Bin: 0.1 (g), 0.5 (j). Zoom: rescaled *y*-axis to visualize the tail of the distribution. Abbreviations: SPT, single particle tracking; SHOT-R, Spatiotemporal High Order Trajectory Reconstruction; *v_i_* and *a_i_*, frame to frame velocity and acceleration; *v_s_* and *a_s_*, instantaneous velocity and acceleration. Data information: **** P < 0.0001. Values are median with interquartile range. Data are not normally distributed (Shapiro-Wilk test), Kruskal–Wallis test followed by Dunn’s multiple comparison. Total number of particle analysed: 372 (b). Each data point corresponds to the percentage of a particle trajectory belonging to that frame to frame velocity category (b).

We finally exploited two unique features of SHOT-R, namely the ability to calculate instantaneous velocity and acceleration (Extended Data Fig. S1b, Fig. 4c,d). SPT reconstruction inherently implies that the particle velocity between two consecutive frames (the smallest temporal resolution) is constant and by extension that its acceleration is equal to zero (Extended Data Fig. S1b, Fig. 4c). On the contrary, SHOT-R does not have such an assumption. Its continuous, high order polynomial function allows, for the first time, to assess instantaneous velocity at any spatiotemporal point of the trajectory including between frames when the exact position of the particle is not captured by a microscope image (Extended Data Fig. S1b). Importantly, since instantaneous velocity can be computed, particle acceleration can also be investigated. Both particle instantaneously velocity and acceleration were best visually represented by plotting their magnitude on top of the particle trajectory (Fig. 4e,h) and heatmap (Fig. 4f,i). When examining the entire dataset, frequency distribution of instantaneous velocities shows a peak between 0.2 and 0.3 *μm/s* with values up to 0.83 *μm/s* (4g, Extended Data Fig. S3e) and acceleration peaks between 0 and 0.5 *μm/s^2^* (4j, Extended Data Fig. S3f). In addition, analysis of axonal pre-miRNA kinematics revealed that maximum velocities (4d, Extended Data Fig. S3e) did not systematically correspond to maximum accelerations (change in velocities) (4i, Extended Data Fig. S3f). To sum up, SHOT-R can analyse particle kinematics globally, on large spatiotemporal scale, like other available tools and it is, however, unique and versatile, since it can also compute particle kinematics locally, at small scale, at high spatiotemporal resolution. Critically, SHOT-R is the only numerical tool available to date to provide insight into particle instantaneous velocities and acceleration.

### 2.5 SHOT-R: advances in reconstructing particle directionality

Most particles display directed movements. They are transported within and between compartments, between the intracellular and extracellular milieus, and/or between cells (Fig. 5a). Cells also exhibit movements in physiological and pathophysiological conditions such as cell chemotaxis during development or metastasis in cancer. Such particle and cellular directionality has important biological implications because it provides insight as to where a specific function is exerted (Fig. 5a). The kinematics study of directionality is, however, strongly limited with current methods. In kymograph-based directionality analysis, a line is manually traced that serves as a reference for the sense of the particle direction along the *x* axis whilst *y* and *z* coordinates are ignored. Kymographs are not able to investigate particle directionality in 3D nor at a spatiotemporal resolution higher than the traced segments (*d_x_/d_t_*, see Fig. 1d, Extended Data Fig. S1a). The SPT algorithms available, on the other hand, do not systematically provide information on particle directionality. To extract this information, individual research groups, therefore, normally rely on in-house scripts, tailored to the specific research question under study, with arbitrary criterias (e.g. directionality defined based on angular displacement, on net linear displacement or on flux value relative to a reference line). These scripts are hitherto not made publically available. Overall, none of the existing approaches allow a standardized analysis in three dimensions (*x, y, z*) and at high temporal resolution. In addition, none integrate at once information on velocity with directionality; directionality is computed *a posteriori* and separately from velocity analysis. SHOT-R, on the other hand, is capable of investigating the kinematics of particle directionality in 3D, at high spatiotemporal resolution, and in an integrated manner during the analysis. Critically, SHOT-R allows, for the first time, to study directionality in a standardized way through a single platform thanks to the possibility of selecting different reference vectors depending on the biological question asked (Fig. 5a,b). The type of direction taken by a particle depends on the type of biological system under study: e.g. anterograde-retrograde, dorsal-ventral, in-out (Fig. 5a). In mathematical terms, this means that a reference vector **d** = (*d_x_, d_y_, d_z_*) is needed and SHOT-R allows to choose between two types of reference to set the particle directionality: Option 1) the vector is a straight line traced along the directionality under study (*x-,y-* or *z*-axis); Option 2) the reference is drawn along any geometrical path of interest (e.g. an axon, or microtubules in a cell etc.) (Fig. 5b). Option 1 is the most time-efficient strategy: the reference vector coincides with a Cartesian axis, and thus it is defined directly in the SHOT-R algorithm without the need of any further input. Differently, for Option 2, *xyz* coordinates to draw a reference are needed (see Methods S1.11).

**Figure 5:**
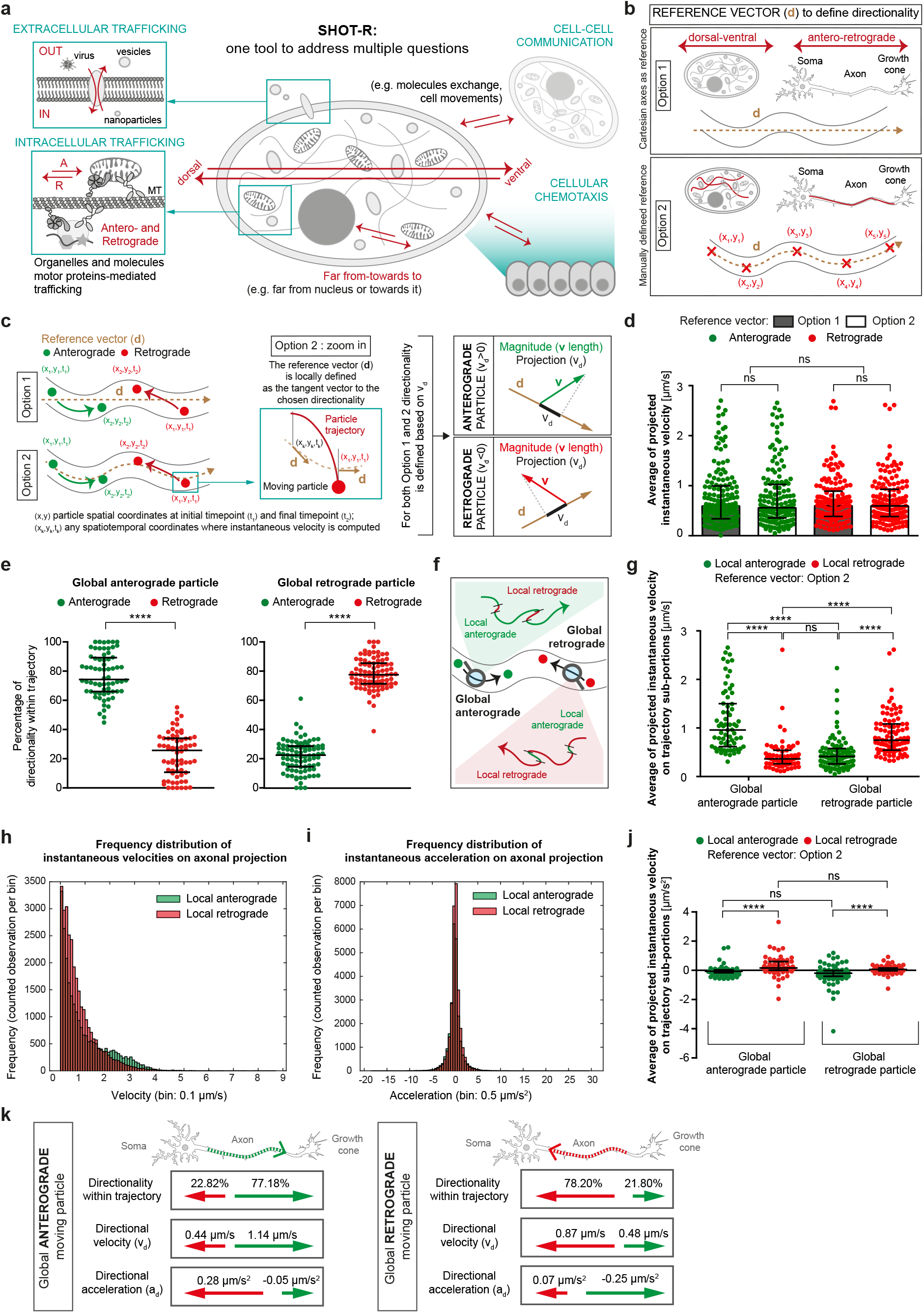
SHOT-R: advances in reconstructing particle directionality. **(a)** Schematic of SHOT-R versatility in particle directionality studies. **(b)** Schematics of reference vector **d**. Option 1: reference along Cartesian axes; Option 2: reference manually defined along any direction of interest. **(c)** Schematic of the computational approach to define particle directionality. **(d)** Average of projected instantaneous velocities using either Option 1 (*x*-axis) or Option 2 (along axon) as reference vector **d**. **(e)** Anterograde and retrograde frequency (in percentage) computed within trajectories of particles globally moving anterogradely (left) or retrogradely (right). **(e)** Schematic defining both global and local particle directionality. **(g)** Average of projected instantaneous velocities for global and local anterograde and retrograde particle movements. **(h,i)** Frequency distribution of the directional instantaneous velocity (*v_d_*, h) and directional instantaneous acceleration (*a_d_*, i). **(j)** Average of projected instantaneous acceleration for global and local anterograde and retrograde particle movements. **(k)** Final schematic of the results. Green and red arrows in the boxes represent respectively local anterograde and retrograde particle directionality. Values reported are means. Data information: ns, not significant; **** P < 0.0001. Each data point corresponds to one particle (d,e,g,j). Values are median with interquartile range. Data are not normally distributed (Shapiro-Wilk normality test), two-tailed Mann-Whitney test (e), Kruskal–Wallis test followed by Dunn’s multiple comparison (d,g,j). Total number of particles analysed: 158 (d,e,g,j) including 68 global anterograde and 90 global retrograde.

Particle directionality can not always be studied along a straight line (e.g. due to the sample shape or a non-linear trajectory) (Fig. 5a,b). Therefore, Option 2, although more time-consuming than Option 1, allows to trace a reference vector along any direction, following the geometry of any sample (Fig. 5a). In physics terms, a particle velocity at time *t* corresponds to a vector **v** (Fig. 5c). Independently of the chosen option, the kinematics of particle directionality is analysed with SHOT-R by projecting the velocity vector **v** onto the reference directionality vector **d**. A vector *v_d_* is thus obtained along the selected direction (Fig. 5c). If *v_d_* > 0, the particle travels towards the direction defined by the vector **d**, whilst if *v_d_* < 0, then it moves in the opposite direction (Fig. 5c).

Here, as an illustrative example, SHOT-R was applied to pre-miRNAs trafficking in axons [5]. Considering the curvilinear axonal shape, both Option 1 and 2 reference vectors were tested on these dataset investigating particle directionality towards the tip of the axons and away from the soma (anterograde transport) or in the opposite direction (retrograde transport) (Fig. 5b,c). When we re-analysed pre-miRNA directionality [5] with SHOT-R, we observed no differences with respect to the selected reference vector in the percentage of global anterograde and retrograde particles per biological replicate (Extended Data Fig. S4a), nor in the average instantaneous velocity (Fig. 5d) and acceleration (Extended Data Fig. S4b). This implies that for this particular dataset Option 1 and 2 are equivalent. Hence, a simple vector along the *x*-axis is a suitable reference for studying anterograde-retrograde movements in axons (Fig. 5b,c, Option 1).

We next analysed change in directionality using our new numerical method. We found that, remarkably, only a few particles displayed a single directionality per trajectory (5 out of 68 anterograde particles moved anterogradely 100% of the time and 3 out of 90 retrograde particles moved retrogradely 100% of the time) (Fig. 5e). We then asked whether changes in directionality within the same trajectory were correlated with a change in instantaneous velocity and/or acceleration. Particles moving anterogradely on sub-portions of the trajectory (thereafter referred to as ”local anterograde particles”, Fig. 5f) were faster when moving globally anterogradely than globally retrogradely (1.14 *μm/s* vs 0.48 *μm/s;* Fig. 5g). When we examined the frequency distribution of instantaneous velocities, we could indeed observe, that the velocity of local anterograde particles peaks both at 0-0.1 *μm/s* and 2.1-2.2 *μm/s* while local retrograde particles peaks only at 0-0.1 *μm/s*; Fig. 5h). This is somewhat expected because the general view of axonal transport is that particles move faster anterogradely due to faster-processing molecular motor kinesins and slower retrogradely due to the slower-processing molecular motor dynein. Intriguingly, however, the velocity of particles didn’t follow this scheme when particles move opposite to the global direction. The velocity of local anterograde and retrograde particles moving opposite to the main direction did not significantly differ (0.48 *μm/s* vs 0.44 *μm/s*; Fig. 5g) and local retrograde particles moving globally retrogradely were faster than local anterograde particles moving globally retrogradely (0.88 *μm/s* vs 0.44 *μm/s*; Fig. 5g). SHOT-R thus reveals that such axonal directed movements are likely more complex than previously envisioned and intricately depends on the global direction, thus the final destination, of the particle. We next examined for the first time particle acceleration. Most particles’ didn’t change velocity with a distribution of particle acceleration peaking at 0-0.5 *μm/s*^2^ (Fig. 5i) and particles moving towards their main direction didn’t change velocity (local anterograde particles moving globally anterogradely: -0.08 *μm/s*^2^, and local retrograde particles moving globally retrogradely 0.07 *μm/s*^2^ (Fig. 5j). However, local anterograde and retrograde particles that moved against their global direction appear to, respectively, decelerate towards the soma (−0.25 *μm/s*^2^) and accelerate towards the growth cone (0.28 *μm/s*^2^) (Fig. 5j). SHOT-R thus enables to reveal novel behaviour whereby axonal RNAs change velocity with direction reversal.

As aforementioned, SHOT-R is unique, since it can also integrate directionality and velocity Eq. (S43). If a particle moves at the same velocity but with different direction at two different time points 1 and 2, with velocity vector 1 pointing closer to the main direction compared to velocity vector 2, then the projection *v*_*d*1_ > *v*_*d*2_ (Extended Data Fig. S4c). In other words, even if **v**_1_=**v**_2_, **v**_1_ contributes more than **v**_2_ to displace the particle along the global direction towards its final destination. Current methods (kymographs and SPT) are strongly limited. Indeed since directionality is computed separately from velocity analysis, these tools assume equal contribution of **v**_1_ and **v**_2_ and risk overestimating particle velocity in a given global direction. To appreciate the impact of such overestimation, we computed, using SHOT-R the directional velocity (*v_d_*) and compared it to the magnitude of velocity vector *v_M_* (used in current methods). We measured that the average instantaneous velocity was lower by 10-22% regardless of the global or local directionality (Extended Data Fig. S4d-f) when *v_d_* was computed compared to *v_M_*. Similarly, frequency distribution also revealed lower instantaneous velocity and accelerations (Extended Data Fig. S4g,h). Taken together, this data suggests that only part of the particle velocity contributes to its displacement along the main direction. Overall, SHOT-R is much better suited than current tools for directionality analysis as it does not overestimate velocity.

By applying SHOT-R on pre-miRNAs particle trafficking, we show the capability of the proposed method to study complex kinematics phenomena from global directionality to change in velocity and acceleration within sub-portions of trajectory (Fig. 5k). This enables to reveal novel biological insight about molecular-motor mediated transport that was hitherto overlooked due to inadequate tools.

### 2.6 SHOT-R: investigating pausing events independently of the acquisition settings

Pausing events are critical aspects of particle transport. Particles are rarely in motion in a continuous manner and instead, their movement can be interrupted with pauses [22, 23, 24, 25]. Such pauses are important in biological terms because they correlate with possible site(s) of function and cargo delivery [25] or direction reversal [22]. In axons, pauses are, for instance, observed when vesicles or cargos interact with organelles [22] or when cargos switch to different microtubules tracks [26, 24]. Both kymographs and SPT can provide information on pausing events (such as existence, frequency, and duration). However, in both approaches, the definition of a pause varies from study to study: first due to the selection or not of motile particles (Step1, Extended Data Fig. S5a), second because pausing events are defined based on parameters set by specific acquisition settings and post-acquisition analysis (i.e. acquisition frequency, time interval and how the velocity is computed) (Step2, Extended Data Fig. S5a). This variability leads to the impossibility of comparing and integrating studies acquired with different settings, slowing down the discovery process despite the many existing studies on particle trafficking.

SHOT-R overcomes this limitation. Indeed, thanks to the continuous reconstruction of the trajectory, it allows the calculation of the *bona fide* instantaneous velocity even between frames (Extended Data Fig. S1b) and thus gives the possibility of defining pausing and other biological events (e.g. reversal, particle motility or immotility) without taking into account acquisition settings. As an example of the use of SHOT-R for this purpose, we applied different thresholds on particle motility (Extended Data Fig. S5b) and different definitions of pausing events [22, 27, 28, 29] (Extended Data Fig. S5e) to our pre-miRNA trafficking dataset [5]. For each test (Extended Data Fig. S5b,e) the percentage of pausing particles and the number of pausing events per particle were computed (Extended Data Fig. S5c,d,f-h). These were significantly decreased when motile particles were taken into account, compared to the entire dataset not filtered for motility (Extended Data Fig. S5b-d) and dramatically different depending on the applied definition of pausing events based on instantaneous velocity (Extended Data Fig. S5e-g). In particular, when pauses were analysed based on instantaneous velocities below 0.1 *μm/s* or 0.2 *μm/s* [28, 27, 29] only ~12-37% of all particles were found to be pausing, while below the 0.4 *μm/s* threshold [22] ~80% of particles were pausing (Extended Data Fig. S5f). Clearly, we can here appraciate that the biological conclusions on the percentage of pausing particles in the dataset depend on how pausing events are defined. The lack of consistency in the results due to different definitions prevents a direct comparison of pausing behaviour between published dataset.

In conclusion, by uncoupling analyses from experimental settings, SHOT-R offers the possibility to reanalyze, compare and integrate the already existing datasets in order to harmonize the knowledge of trafficking phenomena.

### 2.7 SHOT-R: applications on quantitative co-trafficking analysis

The vast majority of cargos are not transported alone within and between cellular compartments but in association with other molecules. For instance, RNA, proteins and organelles are transported docked to molecular motors along microtubule tracks [30, 31, 32, 33, 34]. Co-trafficking analysis is routinely performed to examine through live imaging, whether two or more molecules are co-transported. Although crucial to gain biological insight on mechanisms of molecular transport, this type of investigation is challenging because it requires simultaneous image acquisition. Some expensive instruments have been developed for this purpose for instance spinning disc confocal microscopes. Yet, because of the cost incurred, numerous co-trafficking analyses are still performed with wide field epifluorescence microscopes from single-channel time-lapse acquisition. In other words, the position of two particles are alternately and not simultaneously acquired through the course of a movie. Information on the topographical location of the other particle is thus missing at each acquisition (Fig. 6a).

**Figure 6:**
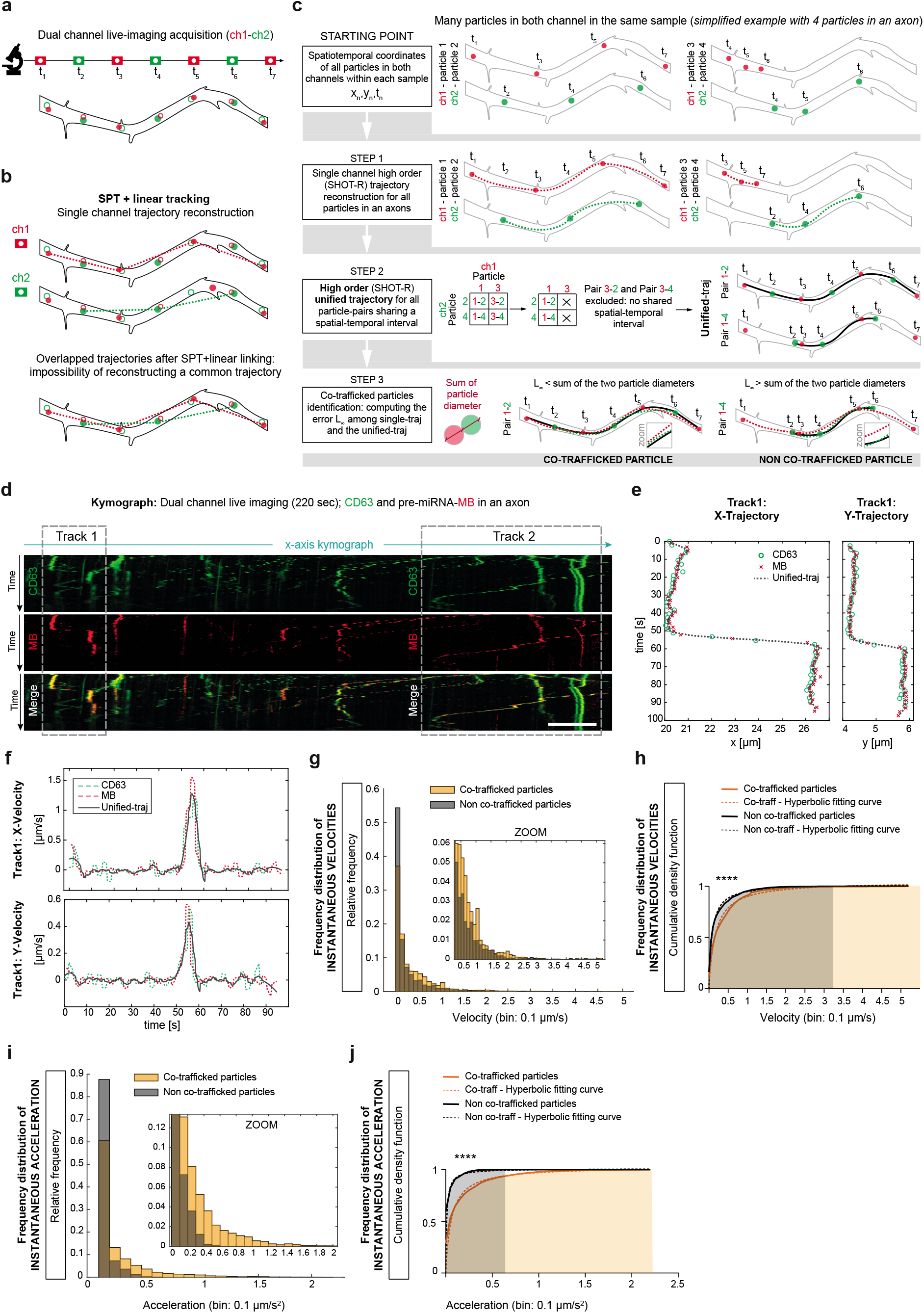
SHOT-R: applications on quantitative co-trafficking analysis. **(a)** Schematic of a not simultaneous dual channel acquisition. **(b)** Schematic of the challenge in reconstructing a unified common trajectory for co-trafficked particles with the current SPT method. **(c)** SHOT-R pipeline to identify co-trafficked particles and reconstruct their unified trajectories. **(d)** Representative kymograph traces of co-trafficked CD63 and pre-miRNA (adapted from [5]). Boxes indicate two representative co-trafficking events (Track1 and 2). **(e)** Unified high order *x*- and *y*-trajectory of Track1 co-trafficked particles. Single green dots and red crosses represent particle coordinates retrieved by TrackMate from raw movies. **(f)** Representative instantaneous velocity plots for co-trafficked particles computed for Track 1. **(g-j)** Frequency distribution of instantaneous velocities (g,h) and acceleration (i,j) show as relative probability (g,i) and cumulative density (h,j). Abbreviations: MB, molecular beacon (recognizing the endogenous pre-miR-181a-1, red); CD63, tetraspanin marker of late endosome/lysosome (green); Unified-traj, unified reference trajectory. Data information: **** P < 0.0001. Data are not normally distributed (Shapiro-Wilk normality test) (g-j). Dashed black and orange lines represent least-squares fit to a hyperbolic equation; fit curve distribution comparison with extra sum-of-squares F test (h,j). 3 independent experiments, 9 axons. Total number of particles analysed: MB-pre-miR-181a-1 (87), CD63 (69), co-trafficked particle-pairs (31) (g-j).

Co-trafficking is usually performed by visual inspection of composite kymograph traces [35, 36, 37, 38, 39, 5] and particles are considered as co-trafficked when their respective traces overlap. Although such an approach succeeds in identifying and counting co-trafficking events, it presents some limitations: i) visual inspection depends on the individual and is prone to biases, ii) manually traced overlapping paths do not allow to accurately describe the single trajectory of the co-transported particles and as a consequence the co-trafficking kinematics remains elusive. The investigation of co-trafficked particle velocity by kymograph would be possible only by automatically identifying a common trace describing the movement of both particles. However, such a tool does not exist yet, and manually tracing this common path would only increase the approximation to this already poorly accurate approach (Fig. 1d,g). In addition, SPT does not address co-trafficking study at all, leaving a methodological gap in all the research fields where kymograph can not be exploited (i.e. when it is not possible to trace a single line over the sample capturing all moving particles within it).

On principle, SPT combined with a reconstruction of a common trajectory of co-trafficked particles would enable the study of co-transported particle velocity. However, this represents a challenge not overcome yet. Indeed, linear linking assumes constant velocity between consecutive frames (Fig. 1e, Extended Data Fig. S1a) and when co-trafficked particles show a non-uniform movement, the reconstruction of a common trajectory through linear linking is simply impossible (as depicted in Fig. 6b). SHOT-R, because of its continuous non-linear trajectory reconstruction, overcomes the limitations and challenges of the current approaches, offering a solution to study for the first time particle co-trafficking kinematics. In particular, we propose a novel strategy to automatically identify co-trafficked particles, reconstruct their common trajectory and from it, assess velocity and acceleration (Fig. 6c). This is achieved through the following steps.

First, the trajectories of all particles within the same axon are reconstructed starting from the spatiotemporal coordinates (**x**_*k*_, *t_k_*) (Fig. 6c, Step1). Second, a high order unified trajectory is built for all particle-pairs sharing a temporal and spatial interval (Fig. 6c Step2, see Methods S1.12 for details). Third, the difference between the single trajectories and this high order unified reference trajectory is computed as *L*_∞_ norm. The *L*_∞_ norm assesses the maximal pointwise error between two curves (Eq.(S39), Fig. 2e). For co-trafficking analysis, we use it to measure the maximal allowed distance between the two single trajectories and the unified one. In SHOT-R, for two particles to be considered as co-trafficked, this distance must be less than the sum of the estimated particle diameter calculated from the single reconstructed trajectory (Fig. 6c, Step3). Ultimately, the co-trafficked particle kinematics is computed for the unified trajectory (Fig. 6c). The sum of the estimated diameters is dataset dependent, thus the *L*_∞_ threshold must be calibrated for each molecules studied.

Thresholding based on *L*_∞_ is extremely stringent considering that a single dot in the trajectory farther than the set threshold is sufficient to exclude a particle pair as non co-trafficked. This strongly limits false positives at the cost of losing some information.

As a proof of concept, we tested our pipeline by examining co-trafficking of pre-miRNA (pre-miR-181a-1, red) with endosome (CD63, in green, a tetraspanin marker of late endosome/lysosome) [5] (Fig. 6d). Contrary to kymograph traces (Fig. 6d), co-trafficked particles are automatically identified and the velocity computed on the reconstructed common trajectory (Fig. 6e, Extended Data Fig. S6a-c), considering both *x* and *y* contribution (Fig. 6f, Extended Data Fig. S6d).

Since the estimated diameter of each particle (pre-miRNA and CD63) was 0.6 *μm* by TrackMate analysis, we therefore set *L*_∞_ ≤ 1.2 μm as a threshold to identify co-trafficking events (Extended Data Fig. S6e). Importantly, the global error *L*_2_ (see Fig. 2e, Eq.(S39) for definition) measured for co-trafficked green-red particles between the common trajectory and the single ones was close to zero (i.e. *L*_2_ mean ~ 0.05 μm; Extended Data Fig. S6f). This remarkably small error suggests that the common trajectory accurately describes the single moving particle of the pair (Extended Data Fig. S6a,c,g). Once a genuine common trajectory is defined, the instantaneous velocity and acceleration can be computed (Fig. 6g-j). Here we found that the frequency distribution of both instantaneous velocity and acceleration differs statistically between co-trafficked or non co-trafficked particles (Fig. 6h,j). In particular, co-trafficked particles compared to the non co-trafficked move faster (Fig. 6g,h; mean ~ 0.37 *μm/s* vs 0.24 *μm/s*) and contrary to non co-trafficked molecules, their velocity changes in time as indicated by instantaneous acceleration greater than zero (Fig. 6i,j; mean ~ 0.18 *μm/s*^2^ vs 0.04 *μm/s*^2^).

In conclusion, we have shown how SHOT-R is able not only to increase the accuracy in kinematic studies compared to all existing methods, but also to apply, for the first time, such an accuracy to investigate cotrafficking events. We foresee that our new tool will contribute in uncovering co-transported mechanisms which have remained hitherto elusive due to lack of adequate algorithms.

## 3 Discussion

We developed SHOT-R, a new analysis tool for reconstructing particle trajectories and studying their kinematics at unprecedented accuracy. Current approaches for particle movements analysis (i.e. kymograph and single particle tracking, SPT) poorly retrieve the original particle trajectory, sacrificing either the spatial and/or temporal resolution. The result of this approximation is the oversimplification of kinematics leading to potential misinterpretation of the biological data, especially in the case of changes in velocities and complex motions. SHOT-R overcomes this limitation and broadens the range of information that can be unraveled on particle kinematics by its innovative approach: the reconstruction of particle trajectories based on high order polynomials in a piecewise continuous manner. Indeed, higher order reconstruction improves the accuracy in computing particle velocities, and, combined with a continuous trajectory reconstruction it allows to study novel kinematics features never investigated before (i.e. instantaneous velocity, acceleration, directional velocity (*v_d_*), kinematics of co-trafficked particles).

As a proof-of-concept, we applied our new numerical method to a published dataset on axonal trafficking [5]. There is a conundrum in the field: how is it possible that molecules become stored for function at specific places within the axon yet display a constant velocity? SHOT-R analysis of instantaneous velocity and acceleration solves this enigma by revealing that particles decelerate against their global direction when they are getting closer to storage points. Remarkably, particles do not decelerate globally (e.g. by slowing down after being trafficked globally from the soma to the growth cone) but following a direction reversal. Particles would, thereby, be preserved at storage points, in their place of function. Overall, SHOT-R reveals that particle velocity is not constant between consecutive frames, as once was thought through the use of classic methods (kymographs and SPT), and that particles display a much more complex trafficking behaviour with critical biological implications. It is tempting to speculate that our observed change of velocity with direction reversal is due to the fact that cargos are coupled to multiple molecular motors at once [40, 41, 42] these molecular motors have different processing ability and directionality thereby exerting tug-of-war forces, i.e. how much they push or pull in a specific direction in concert or alone [43, 44] and this could lead to local change of direction at slower velocity and varied acceleration. In addition, we found that pre-miRNAs move faster and change their velocity more frequently likely when co-trafficked with late endosomes (Fig. 6g-j). This could indicate that these local velocity changes are complex events, involving not only the coordinated activity of multiple molecular motor proteins but also the dynamics of the endosomes themselves (e.g. bidirectional-motility [20], rotational and translational motions [45]). Overall, SHOT-R demonstrated novel complex behaviour of ncRNAs in axons.

Beyond the analysis of axonal transport, we envision that SHOT-R particle kinematics analysis can be extended to numerous fields of research. Our observation of ncRNA axonal trafficking lay important ground for the future study of molecular motor behaviour, and cargo transport in general, which would otherwise not be possible with the current available tools. SHOT-R can finally determine with high accuracy their *bona fide* maximum and minimum instantaneous velocity of molecular motors, and, through directional velocity analysis, reveal the speculated tug-of war behaviour. In addition, SHOT-R acceleration analysis will help unravel complex physiological and pathological dynamics with changes in velocity. For example, SHOT-R can now quantify how much a drug or a therapy slows down cell migration in cancer models, or slows down the entrance of virus particles in a cell. Conversely, SHOT-R can determine how treatments or stimuli speed up movements during axonal degeneration and regeneration after injury, or speed up the entrance of pharmaceutical nanoparticles into a specific target region. SHOT-R is thus uniquely positioned to reveal novel complex biological behaviours for fundamental but also translational research and drug development.

In conclusion, SHOT-R is extremely versatile and can be applied to any live-imaging experiment where the spatiotemporal coordinates can be retrieved and therefore could impact virtually any field requiring particle kinematics analysis.

## 4 Data availability

All the data generated in this manuscript are available from the corresponding authors upon request.

## 5 Code availability

The code is implemented in Matlab programming language, version R2020a. The source code is available from the corresponding authors upon request, and it will be available upon publication at https://gitlab.com/Boscheri/shot-r.

## 6 Author contributions

E.C. and W.B. conceptualized the method and validated it; W.B. and E.C. implemented the software; E.C. performed the statistical analysis and built Figure panels; E.C, W.B. and M.-L.B. wrote the manuscript; M.-L.B. provided funding.

## 7 7 Acknowledgments

M.-L.B. acknowledges financial support from the grants: Marie Curie Career Integration (618969 GUIDANCE-miR), The G. Armenise-Harvard Foundation, MIUR SIR (RBSI144NZ4) and MIUR PRIN 2017 (2017A9MK4R).

## References

[1] Wang, X. & Schwarz, T. L. The mechanism of Ca2+-dependent regulation of kinesin-mediated mitochondrial motility. Cell 136, 163–174 (2009).

[2] Daniele, J. R., Baqri, R. M. & Kunes, S. Analysis of axonal trafficking via a novel live-imaging technique reveals distinct hedgehog transport kinetics. Biology open 6, 714–721 (2017).

[3] Lacovich, V. et al. Tau isoforms imbalance impairs the axonal transport of the amyloid precursor protein in human neurons. Journal of Neuroscience 37, 58–69 (2017).

[4] Fenn, J. D., Johnson, C. M., Peng, J., Jung, P. & Brown, A. Kymograph analysis with high temporal resolution reveals new features of neurofilament transport kinetics. Cytoskeleton 75, 22–41 (2018).

[5] Corradi, E. et al. Axonal precursor mi rna s hitchhike on endosomes and locally regulate the development of neural circuits. The EMBO journal 39, e102513 (2020).

[6] Sergé, A., Bertaux, N., Rigneault, H. & Marguet, D. Dynamic multiple-target tracing to probe spatiotemporal cartography of cell membranes. Nature methods 5, 687–694 (2008).

[7] Jaqaman, K. et al. Robust single-particle tracking in live-cell time-lapse sequences. Nature methods 5, 695–702 (2008).

[8] Cheezum, M. K., Walker, W. F. & Guilford, W. H. Quantitative comparison of algorithms for tracking single fluorescent particles. Biophysical journal 81, 2378–2388 (2001).

[9] Park, H. Y., Buxbaum, A. R. & Singer, R. H. Single mrna tracking in live cells. Methods in enzymology 472, 387–406 (2010).

[10] Meijering, E., Dzyubachyk, O. & Smal, I. Chapter nine-methods for cell and particle tracking. in conn, pm, editor, imaging and spectroscopic analysis of living cells optical and spectroscopic techniques, volume 504 of methods in enzymology (2012).

[11] Chenouard, N. et al. Objective comparison of particle tracking methods. Nature methods 11, 281–289 (2014).

[12] Reid, D. An algorithm for tracking multiple targets. IEEE transactions on Automatic Control 24, 843–854 (1979).

[13] Blackman, S. S. Multiple hypothesis tracking for multiple target tracking. IEEE Aerospace and Electronic Systems Magazine 19, 5–18 (2004).

[14] Dorn, J. F., Danuser, G. & Yang, G. Computational processing and analysis of dynamic fluorescence image data. Methods in cell biology 85, 497–538 (2008).

[15] Meijering, E., Dzyubachyk, O., Smal, I. & van Cappellen, W. A. Tracking in cell and developmental biology. In Seminars in cell & developmental biology, vol. 20, 894–902 (Elsevier, 2009).

[16] Rohr, K. et al. Tracking and quantitative analysis of dynamic movements of cells and particles. Cold Spring Harbor Protocols 2010, pdb–top80 (2010).

[17] Lee, B. H. & Park, H. Y. Hybtrack: A hybrid single particle tracking software using manual and automatic detection of dim signals. Scientific reports 8, 1–7 (2018).

[18] LeVeque, R. Finite Difference Methods for Ordinary and Partial Differential Equations (SIAM, Philadelphia, Pennsylvania, 2007).

[19] Tinevez, J.-Y. et al. TrackMate: An open and extensible platform for single-particle tracking. Methods 115, 80–90 (2017).

[20] Maday, S., Twelvetrees, A. E., Moughamian, A. J. & Holzbaur, E. L. Axonal transport: cargo-specific mechanisms of motility and regulation. Neuron 84, 292–309 (2014).

[21] Leung, K.-M. et al. Cue-polarized transport of *β*-actin mRNA depends on 3’ UTR and microtubules in live growth cones. Frontiers in cellular neuroscience 12, 300 (2018).

[22] Zajac, A. L., Goldman, Y. E., Holzbaur, E. L. & Ostap, E. M. Local cytoskeletal and organelle interactions impact molecular-motor-driven early endosomal trafficking. Current Biology 23, 1173–1180 (2013).

[23] Che, D. L., Chowdary, P. D. & Cui, B. A close look at axonal transport: Cargos slow down when crossing stationary organelles. Neuroscience letters 610, 110–116 (2016).

[24] Yogev, S., Cooper, R., Fetter, R., Horowitz, M. & Shen, K. Microtubule organization determines axonal transport dynamics. Neuron 92, 449–460 (2016).

[25] Guedes-Dias, P. et al. Kinesin-3 responds to local microtubule dynamics to target synaptic cargo delivery to the presynapse. Current Biology 29, 268–282 (2019).

[26] Gu, Y. et al. Rotational dynamics of cargos at pauses during axonal transport. Nature communications 3, 1–8 (2012).

[27] Gluska, S. et al. Rabies virus hijacks and accelerates the p75ntr retrograde axonal transport machinery. PLoS Pathog 10, e1004348 (2014).

[28] Forte, L. A., Gramlich, M. W. & Klyachko, V. A. Activity-dependence of synaptic vesicle dynamics. Journal of Neuroscience 37, 10597–10610 (2017).

[29] Yardeni, T. et al. High content image analysis reveals function of mir-124 upstream of vimentin in regulating motor neuron mitochondria. Scientific reports 8, 1–13 (2018).

[30] Ross, J. L., Ali, M. Y. & Warshaw, D. M. Cargo transport: molecular motors navigate a complex cytoskeleton. Current opinion in cell biology 20, 41–47 (2008).

[31] Barlan, K. & Gelfand, V. I. Microtubule-based transport and the distribution, tethering, and organization of organelles. Cold Spring Harbor perspectives in biology 9, a025817 (2017).

[32] Martin, K. C. & Ephrussi, A. mrna localization: gene expression in the spatial dimension. Cell 136, 719–730 (2009).

[33] Dalla Costa, I. et al. The functional organization of axonal mrna transport and translation. Nature Reviews Neuroscience 22, 77–91 (2021).

[34] Zerial, M. & McBride, H. Rab proteins as membrane organizers. Nature reviews Molecular cell biology 2, 107–117 (2001).

[35] Salogiannis, J., Egan, M. J. & Reck-Peterson, S. L. Peroxisomes move by hitchhiking on early endosomes using the novel linker protein pxda. Journal of Cell Biology 212, 289–296 (2016).

[36] Bowen, A. B., Bourke, A. M., Hiester, B. G., Hanus, C. & Kennedy, M. J. Golgi-independent secretory trafficking through recycling endosomes in neuronal dendrites and spines. Elife 6, e27362 (2017).

[37] Gershoni-Emek, N. et al. Localization of rnai machinery to axonal branch points and growth cones is facilitated by mitochondria and is disrupted in als. Frontiers in molecular neuroscience 11, 311 (2018).

[38] Cioni, J.-M. et al. Late endosomes act as mRNA translation platforms and sustain mitochondria in axons. Cell 176, 56–72 (2019).

[39] Liao, Y.-C. et al. Rna granules hitchhike on lysosomes for long-distance transport, using annexin a11 as a molecular tether. Cell 179, 147–164 (2019).

[40] Ashkin, A., Schütze, K., Dziedzic, J., Euteneuer, U. & Schliwa, M. Force generation of organelle transport measured in vivo by an infrared laser trap. Nature 348, 346–348 (1990).

[41] Gross, S. P., Welte, M. A., Block, S. M. & Wieschaus, E. F. Coordination of opposite-polarity microtubule motors. The Journal of cell biology 156, 715–724 (2002).

[42] Kural, C. et al. Kinesin and dynein move a peroxisome in vivo: a tug-of-war or coordinated movement? Science 308, 1469–1472 (2005).

[43] Lipowsky, R., Beeg, J., Dimova, R., Klumpp, S. & Muller, M. J. Cooperative behavior of molecular motors: cargo transport and traffic phenomena. Physica E: Low-dimensional Systems and Nanostructures 42, 649–661 (2010).

[44] Klumpp, S., Keller, C., Berger, F. & Lipowsky, R. Molecular motors: Cooperative phenomena of multiple molecular motors. In Multiscale Modeling in Biomechanics and Mechanobiology, 27–61 (Springer, 2015).

[45] Kaplan, L., Ierokomos, A., Chowdary, P., Bryant, Z. & Cui, B. Rotation of endosomes demonstrates coordination of molecular motors during axonal transport. Science Advances 4, e1602170 (2018).

[46] Stroud, A. Approximate Calculation of Multiple Integrals (Prentice-Hall Inc., Englewood Cliffs, New Jersey, 1971).

