## Supplementary material for "SHOT-R: A next generation algorithm for particle kinematics analysis"

### S1 Methods

#### S1.1 Particle coordinates

In our biological applications, pre-miRNAs particle $xy$ coordinates are extracted from published movies [5] with TrackMate v5.2.0 [19]. Raw movies are cropped the axonal portion to analyse. The leading tip of the axon, the growth cone, is kept on the right side of each crop, which is not included in the analysis since the high density in particle signal does not allow to recognize single pre-miRNA puncta. Occasionally, background correction with rolling ball pixel size 10 is applied to increase signal to noise ratio in ImageJ. Difference of Gaussian particle detection (DoG) is used as detector in TrackMate. The estimated diameter is set to 0.6 $\mu m$ and the threshold in the TrackMate detection step is fixed from 10 to 400 depending on the signal to noise ratio of each movie. Median filter, reducing the generation of spurious spots in case of salt and pepper noise, and sub-pixel localization options are selected. The tracking settings are: link max distance "1"; gap closing max distance "2" and gap closing max frame gap "4" (for movies with single channel acquisition); link max distance "5"; gap closing max distance "2" and gap closing max frame gap "1" (for movies with both green and red channel). Splitting or merging events are not expected, hence Simple Linear Assignment Problem (LAP) Tracker is used. No filters are applied to the track step. The automatically tracked lines are manually checked and minor adjustments applied (i.e. remove signal coming from other axons present in the same field; link tracks belonging to the same particle, only when the particle is visible but not detected due to the applied threshold). Prior to analysis a filter on number of consecutive frames per track is applied in order to clear spurious links done by the automatic tracking. For movies on pre-miRNA endogenous trafficking the minimal number of frames is set at 20, for co-trafficking movies at 5.

#### S1.2 Split trajectories

Once the space-time coordinates $(\mathbf{x}_k, t_k)$ are given for each data point $k = 1, \ldots, N_K$ (Fig. 2a), we proceed in a dimension-by-dimension manner by splitting the position vector $\mathbf{x} = (x, y, z)$ along each spatial direction (Fig. 2b). Specifically, the issue related to a multidimensional spatial domain is tackled by considering the trajectory equations

$$\begin{cases} \frac{dx}{dt} = v_x(x,t), & x(t=0) = x_0, \\ \frac{dy}{dt} = v_y(y,t), & y(t=0) = y_0, \\ \frac{dz}{dt} = v_z(z,t), & z(t=0) = z_0, \end{cases} \tag{S1}$$

that directly derive from the original definition (1). This strategy simplifies the mathematical approach and permits to save computational time while keeping the desired temporal and spatial resolution of fully 3D procedures. Let observe that all equations in (S1) are mathematically the same object, therefore the novel SHOT-R method is based on the solution of a generic one-dimensional trajectory of particle along the direction $s$, i.e.

$$\frac{ds}{dt} = v_s(s,t), \quad s(t=0) = s_0, \tag{S2}$$

where $s$ can be referred to any spatial direction among $\mathbf{x}$ and the same holds true for the velocity component $v_s$ with $\mathbf{v}$. This makes our algorithm very general and applicable to a wide range of physical and biological phenomena. We stress that the only information needed as input are the point coordinates $\mathbf{x}_k$ defined along the trajectory that has to be reconstructed.

Let $\mathcal{D}$ represent the spatial dimensionality of the problem under consideration, i.e. $\mathcal{D} \in [1,3]$. Consequently, $\mathbf{x}_k \in \mathbb{R}^{\mathcal{D}}$, that is the spatial coordinates are given by real numbers in dimension $\mathcal{D}$. Along each spatial direction we have then the set of space-time coordinates $(s_k, t_k)$ that are extracted from $(\mathbf{x}_k, t_k)$. What carried out for the generic direction $s$ will be performed as many times as the dimensionality of the problem is matched, thus a genuinely multidimensional trajectory defined in dimension $\mathcal{D}$ is reduced to a total number of $\mathcal{D}$ one-dimensional trajectories. The full multidimensional trajectory can then be simply gathered back by collecting all one-dimensional contributions.

#### S1.3 Computational mesh

The time coordinate $t$ is defined in the time interval $T = [t_1; t_{N_K}]$, i.e. $t \in T$, while the space coordinate $s$ is bounded in $\Omega = [s_1; s_{N_K}]$, i.e. $s \in \Omega$. The computational domain $\Omega$ is discretized with a total number of $N_I = N_K - 1$ cells with size $\Delta t_i$ and barycenter time coordinate $t_i$ given by

$$\Delta t_i = t_{i+1/2} - t_{i-1/2}, \qquad t_i = \frac{t_{i+1/2} + t_{i-1/2}}{2}, \qquad \text{for} \quad i = 1, \dots, N_I, \tag{S3}$$

where the values at the interfaces are directly available from $t_{i-1/2} = t_k$ and $t_{i+1/2} = t_{k+1}$ (Fig. 2c). Let observe that index $k$ is used for cell interfaces $i - 1/2$ in order to obtain a compatible notation on the computational mesh, thus one has the equivalence $k = i - 1/2$. Notice that we do not assume equidistant points, thus the trajectory $s(t)$ can be defined by a sequence of space-time coordinates $(s_k, t_k)$ that are not regularly extracted in time. This does not cause any problem in the algorithm since each cell is assigned its own width $\Delta t_i$. The resulting computational mesh is referred to as staggered mesh, because the known quantities are defined at the interfaces $(k, k+1)$ of each cell $i$ whereas the sought unknown reconstruction polynomial $p_i^N(s,t)$ will be discretized at the cell center $s_i$. In other words, known data and unknown reconstruction polynomials adopt staggered definitions on the computational mesh.

#### S1.4 Reconstruction algorithm

The aim of the reconstruction procedure is to obtain a piecewise high order polynomial approximation of the particle trajectory. The result is then a set of polynomials $p_i^N(s,t)$ of degree $N \geq 1$ that are defined within each computational cell $i$ for each spatial direction $s$. The polynomial of arbitrary degree $N$ is written in terms of a normalized Taylor series expanded around the barycenter coordinate $t_i$, that is

$$p_i^N(s,t) = \sum_{q=0}^{N} \frac{(t - t_i)^q}{q!\, \Delta t_i^q}\, \hat{s}_i := \phi_l(t)\, \hat{s}_{i,l}, \tag{S4}$$

with $\Delta t_i^q$ being a normalizing factor necessary to avoid ill-conditioned reconstruction matrices and $q!$ represents as usual the factorial of $q$. The unknown degrees of freedom are denoted by $\hat{s}_i$ which have to be determined by the reconstruction algorithm. Let observe that the polynomial (S4) can also be written as an expansion with a set of basis functions $\phi_l(t) \in \mathbb{P}^N$ belonging to the space of polynomials of degree $N$, i.e. $\mathbb{P}^N$. Einstein summation convention is adopted implying summation over repeated indexes, thus the reconstruction polynomial for cell $i$ compactly writes as $\phi_l(t)\, \hat{s}_{i,l}$.

To retrieve a polynomial of degree $N$, a total number of $N+1$ unknowns $\hat{s}_i$ must be uniquely determined, as follows from the definition (S4). Depending on the chosen degree $N \geq 1$, the piecewise high order polynomial is reconstructed for each cell $i$ by considering the surrounding known points, that form the so-called reconstruction stencil $\mathcal{N}_i = \{...i-3/2, i-1/2, i+1/2, i+3/2...\}$ (Fig. 2d). The stencil counts a total number of $2(N+1)$ cell interfaces for even and odd degree $N$, so that the stencil is always symmetric with respect to the cell $i$ under consideration. We address with $N$ the polynomial degree, while $\mathcal{O}(N) = N + 1$ is the order of accuracy of the method. For instance, a polynomial $P1$ of degree $N = 1$ is second order accurate, i.e. $\mathcal{O}(1) = 2$, because it is exact up to degree 1 and the approximation errors arise starting from $N = 2$, thus the accuracy is of second order. The safety factor 2 in the total number of cells contained in the stencil allows for including linear constraints in the reconstruction polynomials, i.e. overdetermined linear systems are assembled. Indeed, we also want to ensure continuity of the reconstructed polynomials across cell interfaces located at $i \pm 1/2$. This would generate a continuous profile for the trajectory, while keeping the reconstruction algorithm local to each cell and therefore computationally efficient.

The reconstruction is designed in such a way that the trajectory coordinates $s_k = s_{i\pm1/2}$ are exactly matched by the polynomial $p_i^N(s,t)$. For each cell $i$ the reconstruction system is constructed as follows:

$$\phi_l(t_k)\, \hat{s}_{i,l} = s_k, \qquad k \in \mathcal{N}_i, \tag{S5}$$

with the linear constraints

$$\phi_l(t_{i-1/2})\,\hat{s}_{i,l} = s_{i-1/2}, \qquad \phi_l(t_{i+1/2})\,\hat{s}_{i,l} = s_{i+1/2}. \tag{S6}$$

The linear reconstruction system (S5)-(S6) set up above can be written in matrix-vector form as

$$\left\{ \begin{array}{l} \mathbf{M}\,\mathbf{S} = \mathbf{B} \\ \mathbf{C}\,\mathbf{S} = \mathbf{D}\,\mathbf{B} \end{array} \right. , \tag{S7}$$

with the definitions

$$\mathbf{B}_{[\mathcal{S}\times 1]} = \begin{bmatrix} \vdots \\ s_{k-1} \\ s_k \\ s_{k+1} \\ \vdots \end{bmatrix}, \qquad \mathbf{S}_{[(N+1)\times 1]} = \begin{bmatrix} \hat{s}_{i,1} \\ \vdots \\ \hat{s}_{i,N+1} \end{bmatrix}, \qquad \mathbf{M}_{[\mathcal{S}\times(N+1)]} = \begin{bmatrix} \phi_1(t_1) & \dots & \phi_{N+1}(t_1) \\ \vdots & \vdots & \vdots \\ \phi_1(t_k) & \dots & \phi_{N+1}(t_k) \\ \vdots & \vdots & \vdots \\ \phi_1(t_{\mathcal{S}}) & \dots & \phi_{N+1}(t_{\mathcal{S}}) \end{bmatrix}. \tag{S8}$$

The vector of unknown coefficients is $\mathbf{S}$, the right hand side which contains the known position values is $\mathbf{B}$ and the reconstruction matrix is given by $\mathbf{M}$. The constrained matrix $\mathbf{C}$ in (S7) is a sub-partition of $\mathbf{M}$ and contains those rows which correspond to the stencil elements $i \pm 1/2$, while the vector $\mathbf{D}$ is zero everywhere apart from the two entries related to the constraints (S6), where it is set to 1. The constrained least squares (CLSQ) method is adopted for solving system (S7), therefore a functional $g(\mathbf{S})$ is defined and minimized by requiring its derivatives to vanish, thus

$$g(\mathbf{S}) = (\mathbf{MW}-\mathbf{B})^T(\mathbf{MW}-\mathbf{B}) - \boldsymbol{\mu}^T(\mathbf{CW}-\mathbf{DB}), \tag{S9}$$

$$\frac{\partial g}{\partial \mathbf{S}} = 2\mathbf{MMS} - 2\mathbf{MB} - \mathbf{C}\boldsymbol{\mu} = 0, \tag{S10}$$

$$\frac{\partial g}{\partial \boldsymbol{\mu}} = -(\mathbf{CW}-\mathbf{DB}) = 0, \tag{S11}$$

where $\boldsymbol{\mu}$ is the vector of Lagrange multipliers to enforce the linear constraints. The associated enlarged linear system of normal equations then reads

$$\begin{pmatrix} 2\mathbf{MM} & -\mathbf{C} \\ \mathbf{C} & \mathbf{0} \end{pmatrix} \begin{pmatrix} \mathbf{S} \\ \boldsymbol{\mu} \end{pmatrix} = \begin{pmatrix} 2\mathbf{M} \\ \mathbf{D} \end{pmatrix} \mathbf{B}, \tag{S12}$$

whose solution can be simply obtained as

$$\begin{pmatrix} \mathbf{S} \\ \boldsymbol{\mu} \end{pmatrix} = \begin{pmatrix} 2\mathbf{MM} & -\mathbf{C} \\ \mathbf{C} & \mathbf{0} \end{pmatrix}^{-1} \begin{pmatrix} 2\mathbf{M} \\ \mathbf{D} \end{pmatrix} \mathbf{B} := \mathbf{R}^L\,\mathbf{B}. \tag{S13}$$

We are only interested in the solution of the expansion coefficients $\mathbf{S}$, therefore the final reconstruction matrix $\mathbf{R}$ for the trajectory along the $s$ dimension is a subset of the CLSQ matrix $\mathbf{R}^L$, that is $\mathbf{R} := \mathbf{R}^L_{[(N+1)\times\mathcal{N}_i]}$. Once the reconstruction matrices have been computed for all cells the reconstruction algorithm is is efficiently carried out by one matrix-vector product, namely

$$\mathbf{S} = \mathbf{R}^L_{[(N+1)\times\mathcal{N}_i]}\,\mathbf{B}, \tag{S14}$$

which permits to obtain the sought degrees of freedom $\hat{s}_{i,l}$ and thus to uniquely define the reconstruction polynomial $p_i^N(s,t)$ according to (S4) within each cell $i$.

#### S1.5 Computation of the length of curvilinear trajectory

The evaluation of the length $L$ of the particle trajectory is needed in order to compute average quantities over the entire path (Extended Data Fig. S3a). The length is defined as the integral along the line drawn by the particle trajectory. The integral can be rewritten as the sum of the integrals over all computational cells $N_I$, that is

$$L = \int_{\mathbf{x}_1}^{\mathbf{x}_{N_K}} dl = \sum_{i=1}^{N_I} \int_{\mathbf{x}_{i-1/2}}^{\mathbf{x}_{i+1/2}} dl. \tag{S15}$$

The coordinates $(\mathbf{x}_1, \mathbf{x}_{N_K})$ denote the starting and the ending point of the trajectory in the space of dimension $\mathcal{D} \in [1;3]$. To perform the integration (S15) at high order of accuracy, it is not sufficient to split the trajectory into $N_I$ segments according to the number of cells and to sum up all the linear contributions. Indeed, this would lead to a second order accurate evaluation of $L$, since piecewise linear polynomials $P1$ are used.

To obtain a consistent high order computation of the trajectory length, isoparametric discretizations must be adopted along the generic spatial direction $s$. To this aim, let us first define a reference coordinate system $\xi \in [0;1]$ which is used to map a generic cell $i$, hence

$$s = s_{i-1/2} + \xi \left( s_{i+1/2} - s_{i-1/2} \right), \tag{S16}$$

so that for $\xi = 0$ the left interface coordinate $s_{i-1/2}$ of the cell is retrieved, while for $\xi = 1$ the right interface $s_{i+1/2}$ is obtained. In the reference system we then define a set of nodal basis functions $\theta_m(\xi)$ which are chosen to be the Lagrange polynomials passing through the set of nodes $\hat{\xi}_m = m/N$ with $0 \leq m \leq N$. Furthermore, the standard interpolation condition $\theta_q(\hat{\xi}_m) = \delta_{qm}$ holds for the basis, with $\delta_{qm}$ being the usual Kronecker symbol. Up to degree $N = 3$ the nodal basis functions $\theta_m(\xi)$ with associated nodes $\hat{\xi}_m$ explicitly write as follows:

- $N = 1$

$$\begin{array}{ll} \theta_1 = 1 - \xi & \theta_2 = \xi \\ \\ \hat{\xi}_1 = 0 & \hat{\xi}_2 = 1 \end{array} \tag{S17}$$

- $N = 2$

$$\begin{array}{lll} \theta_1 = 1 - 3\xi + 2\xi^2 & \theta_2 = 4\xi - 4\xi^2 & \theta_3 = -\xi + 2\xi^2 \\ \\ \hat{\xi}_1 = 0 & \hat{\xi}_2 = 1/2 & \hat{\xi}_3 = 1 \end{array} \tag{S18}$$

- $N = 3$

$$\begin{array}{ll} \theta_1 = 1 - \frac{11}{2}\xi + 9\xi^2 - \frac{9}{2}\xi^3 & \theta_2 = 9\xi - \frac{45}{2}\xi^2 + \frac{27}{2}\xi^3 \\ \theta_3 = -\frac{9}{2}\xi + 18\xi^2 - \frac{27}{2}\xi^3 & \theta_4 = \xi - \frac{9}{2}\xi^2 + \frac{9}{2}\xi^3 \\ \\ \hat{\xi}_1 = 0 & \hat{\xi}_2 = 1/3 \\ \hat{\xi}_3 = 2/3 & \hat{\xi}_4 = 1 \end{array} \tag{S19}$$

The integral over cell $i$ in (S15) can be performed in the reference system at the aid of a change of variables $\mathbf{x} \to \xi$, hence obtaining

$$\int_{\mathbf{x}_{i-1/2}}^{\mathbf{x}_{i+1/2}} dl = \int_0^1 |J_\xi| \, d\xi, \tag{S20}$$

with the Jacobian determinant given by

$$|J_\xi| = \sqrt{\left(\frac{\partial x}{\partial \xi}\right)^2 + \left(\frac{\partial y}{\partial \xi}\right)^2 + \left(\frac{\partial z}{\partial \xi}\right)^2}. \tag{S21}$$

The computation of the derivatives in (S21) is done as usual by exploiting the expansion of the basis functions $\theta(\xi)$, thus for a generic spatial coordinate $s$ it holds

$$\frac{\partial s}{\partial \xi} = \frac{\partial \theta_m(\xi)}{\partial \xi} \hat{s}_m, \tag{S22}$$

where the degrees of freedom $\hat{s}_m$ represent the value of the coordinate $s$ corresponding to the reference node $\hat{\xi}_m$. To be precise, the values $\hat{s}_m$ are computed according to (S23), relying on the SHOT-R reconstruction polynomial. Finally, the integration in (S20) is numerically approximated by Gaussian quadrature formulae of suitable order of accuracy [46].

#### S1.6 Computation of position, velocity and acceleration

The reconstruction polynomials $p_i^N(s,t)$ provide a piecewise continuous representation of the trajectory within each cell $i$ along the generic spatial direction $s$ as a function of time $t$. Thus, the kinematic description (position $s$, velocity $v_s$ and acceleration $a_s$) of the particle at any given time $\tilde{t}$ can be easily computed in two steps:

1. find the computational cell containing $\tilde{t}$, i.e. $t_{i-1/2} \leq \tilde{t} \leq t_{i+1/2}$, so that the associated reconstruction polynomial $p_i^N(s,t)$ can be identified;
2. extract the kinematic quantities

$$s(\tilde{t}) = p_i^N(s,\tilde{t}) = \sum_{q=0}^{N} \frac{(\tilde{t}-t_i)^q}{q!\,\Delta t_i^q}\,\hat{s}_i, \tag{S23}$$

$$v_s(\tilde{t}) = \frac{d\,p_i^N(s,\tilde{t})}{dt} = \sum_{q=0}^{N} \frac{q(\tilde{t}-t_i)^{q-1}}{q!\,\Delta t_i^q}\,\hat{s}_i, \tag{S24}$$

$$a_s(\tilde{t}) = \frac{d^2\,p_i^N(s,\tilde{t})}{dt^2} = \sum_{q=0}^{N} \frac{q(q-1)(\tilde{t}-t_i)^{q-2}}{q!\,\Delta t_i^q}\,\hat{s}_i. \tag{S25}$$

The above definitions are referred to as instantaneous quantities. We also need to give explicit formulation for the global average speed $v_L$, the displacement velocity $v_D$ and the mean of instantaneous velocities within each frame $\bar{v}_M$. These definitions are mathematically described as follows:

$$v_L = \frac{L}{t_{N_K}-t_1}, \qquad L = \int_{\mathbf{x}_1}^{\mathbf{x}_{N_K}} dl, \tag{S26}$$

$$v_D = \frac{s_{N_K}-s_1}{t_{N_K}-t_1}, \tag{S27}$$

$$v_M = \frac{1}{N_I}\sum_{i=1}^{N_I} v_i, \qquad v_i = \frac{s_{i+1}-s_i}{t_{i+1}-t_i}. \tag{S28}$$

with $s$ denoting as usual a generic spatial direction among $\mathbf{x}=(x,y,z)$. The length of the trajectory $L$ in (S26) is computed as the integral along the curvilinear path from $\mathbf{x}_1$ to $\mathbf{x}_{N_K}$ according to (S15). Finally, the instantaneous velocity at a generic space-time coordinate $(s,\tilde{t})$ can be simply evaluated relying on the velocity reconstruction polynomial, thus according to (S24).

In order to fully exploit the piecewise continuous information provided by the high order reconstruction, the evaluation of all instantaneous kinematic quantities (position, velocity and acceleration) is extracted on a very fine computational mesh given by $N+1$ Gauss points leading to exact integration up to order $2(N+1)-1$ (see [46]) defined within each cell. This is done for the analysis reported in Figures 1-5 and it guarantees that the high order polynomials are integrated exactly. For the co-trafficking analysis shown in Figure 6, we always used a total number of 100 Gauss points for each trajectory.

#### S1.7 Numerical convergence studies

In order to validate the accuracy of the proposed reconstruction algorithm, the numerical convergence of the method must be rigorously analysed. The order of accuracy $p>0$ of a numerical method is also defined as the largest value of $p$ for which the following inequality holds

$$\varepsilon(\Delta t) \leq C\,\Delta t^p, \tag{S29}$$

where $\varepsilon(\Delta t)$ is the error of the numerical solution, which depends on the mesh size $\Delta t$. The constant $C$ is independent of $\Delta t$, this is also written thus

$$\varepsilon = \mathcal{O}(\Delta t^p), \qquad \text{as} \quad \Delta t \to 0. \tag{S30}$$

In other words, the numerical error must decrease with convergence rate of $\mathcal{O}(\Delta t^p)$ as the mesh size decreases. Given two regular meshes with mesh sizes $\Delta t_1$ and $\Delta t_2$, respectively, and the same numerical method, we have a fixed relationship between the meshes which writes

$$\varepsilon(\Delta t_1) \;=\; C\Delta t_1^p, \tag{S31}$$

$$\varepsilon(\Delta t_2) \;=\; C\Delta t_2^p. \tag{S32}$$

Dividing (S32) by (S31) we obtain

$$\left(\frac{\Delta t_2}{\Delta t_1}\right)^p = \frac{\varepsilon(\Delta t_2)}{\varepsilon(\Delta t_1)}, \tag{S33}$$

which leads to

$$p = \frac{\ln\left(\frac{\varepsilon(\Delta t_2)}{\varepsilon(\Delta t_1)}\right)}{\ln\left(\frac{\Delta t_2}{\Delta t_1}\right)}, \tag{S34}$$

that is the empirical order of accuracy. It must be $p = N + 1$, meaning that a numerical scheme with reconstructions of degree $N$ introduces numerical errors starting from degree $N + 1$, that is a polynomial of degree $N$ can can exactly reconstructed. In practice one performs numerical experiments for a sequence of refined meshes $\{\Delta t_1, \Delta t_2, \dots, \Delta t_K\}$ with usually a fixed ratio between successive meshes of sizes $\Delta t_k$ and $\Delta t_{k+1}$. For example $\Delta t_{k+1} = \frac{\Delta t_k}{2}$. A sequence of numbers $\{p_2, p_3, \dots, p_K\}$ is then retrieved which is given by

$$p_k = \frac{\ln\left(\frac{\varepsilon(\Delta t_k)}{\varepsilon(\Delta t_{k-1})}\right)}{\ln\left(\frac{\Delta t_k}{\Delta t_{k-1}}\right)}, \qquad k = 2, \dots, K. \tag{S35}$$

To carry out the convergence studies, we consider the time interval $T = [-1; 1]$ and we prescribe the following analytical trajectory in 3D:

$$\mathbf{x} = (\sin(\pi x)\cos(2\pi x),\ 3\cos(2\pi y) - 2\sin(\pi y),\ -6\sin(\pi z) + 2\cos(3\pi z)). \tag{S36}$$

SHOT-R method is then employed to reconstruct the trajectory along each direction and the errors $\varepsilon$ are measured for each spatial direction $(x, y, z)$ in $L_1$, $L_2$ and $L_\infty$ norms according to (S39). The exact solution is explicitly computed with (S36), while the reconstructed position is evaluated according to (S23). A sequence of four successively refined meshes is used with $\Delta t = \{100, 200, 400, 800\}$ and the results are reported in Table S1, where the formal order of accuracy is perfectly achieved by SHOT-R for $N \in [1; 5]$, thus up to sixth order of accuracy. This numerically confirms that the SHOT-R algorithm is actually high order accurate for an arbitrary polynomial degree $N$.

#### S1.8 Mathematical test for SHOT-R accuracy

In order to theoretically demonstrate the capabilities of SHOT-R algorithm, let us consider a time computational domain $T = [t_0; t_f] = [0; 2]$ and a two-dimensional $xy$ space. We prescribe the particle velocity given by

$$v_x(t) = 1 + \tanh(5(t-1)), \qquad v_y(t) = 2\pi\,\cos(2\pi\,t), \tag{S37}$$

from which it is possible to evaluate an analytical expression for the particle trajectory by integrating (S37) over time $t$, thus

$$x(t) = \int_t v_x(t)\,dt = 2t - \frac{\log(\tanh(5(t-1)) + 1)}{5}, \qquad y(t) = \int_t v_y(t)\,dt = \sin(2\pi\,t). \tag{S38}$$

Expressions (S37)-(S38) describe the kinematics of a particle in the temporal domain $T$, which is discretized with three different computational meshes of characteristic mesh size of $\Delta t_1 = 1/10$, $\Delta t_2 = 1/20$ and $\Delta t_3 = 1/40$. In other words, a total number of $N_1 = 21$, $N_2 = 41$ and $N_3 = 81$ equidistant points of coordinates $\mathbf{x}_k = (x_k, y_k)$ are prescribed according to (S38), which constitute the initial condition (IC) of the ODE (1).

#### S1.9 Computing error norms

To quantitatively appreciate the accuracy of the results, the errors between the reconstructed and a reference solution for both position and velocity are measured in $L_1$, $L_2$ and $L_\infty$ norms, that for a generic spatial direction $s$ are

$$L_1 = \int_\Omega \|s_e(t) - s_h(t)\|\, dt, \qquad L_2 = \sqrt{\int_\Omega \|s_e(t) - s_h(t)\|^2\, dt}, \qquad L_\infty = \max_\Omega \|s_e(t) - s_h(t)\|\,. \tag{S39}$$

The reference solution $s_e$ depends on the specific application. In particular, it is given by (S38) and (S37) for position and velocity (Fig. 1), the $P3$ polynomial (Fig. 2) or the unified trajectory (Fig. 6). Then, $s_h(t)$ denotes the numerical solution computed with SHOT-R (or the linear reconstruction in Fig. 2). The integrals appearing in (S39) are evaluated using Gaussian quadrature formulae of suitable accuracy (see [46]).

#### S1.10 Backward integration of particle trajectory

The results shown in Fig. 2h have been obtained by integrating backward in time the trajectory of the particle using the novel SHOT-R strategy. From the mathematical viewpoint, this corresponds to the solution of the ODE

$$\frac{d\mathbf{x}}{dt} = -\mathbf{v}(\mathbf{x}, t), \qquad \mathbf{x}_0 = \mathbf{x}_{N_K}, \qquad t \in [0; t_{N_K} - t_1], \tag{S40}$$

with $\mathcal{D} = 2$ and therefore $\mathbf{x} = (x, y)$ and $\mathbf{v} = (v_x, v_y)$. The initial condition $\mathbf{x}_0$ of the ODE is given by the $xy$ position of the particle at the end of the trajectory, which coincides with the last input point $N_K$. We proceed as follows. Firstly, SHOT-R method is employed for reconstructing the particle trajectory, i.e. for obtaining $p_i^N(\mathbf{x}, t)$ for all cells $i$. Secondly, the velocity vector $\mathbf{v}_i$ is computed with (S24), hence obtaining a high order velocity field defined within each computational cell. Finally, equation (S40) is solved up to the final time $t_f = t_{N_K} - t_1$ in such a way that the position of the particle at the beginning of the trajectory is retrieved. Obviously, due to numerical approximation, this will never coincide with the input coordinates of the first point $\mathbf{x}_1$. However, we have shown that higher order reconstructions provide a more accurate results, that is the final reconstructed coordinates which arise from the solution of the ODE (S40) are closer to the original first point of the trajectory taken as input for the trajectory reconstruction.

In order to solve equation (S40) any classical ODE integrator can be used, provided that it satisfies at least the same accuracy property of the polynomial reconstruction. Here, we rely on a very popular numerical technique, namely the class of Runge-Kutta (RK) schemes [18]. For $P1$ reconstruction (SPT) we use a second order RK method, while for $P3$ (SHOT-R) the fourth order version is adopted. Runge-Kutta schemes explicitly write as follows:

- second order Runge-Kutta (RK2)

$$\begin{aligned} \mathbf{x}_{n+1} &= \mathbf{x}_n + \frac{d\tau}{2}\left(\mathbf{k}_1 + \mathbf{k}_2\right), \\ \mathbf{k}_1 &= -\mathbf{v}(\tau_n, \mathbf{x}_n), \\ \mathbf{k}_2 &= -\mathbf{v}\left(\mathbf{x}_n + \frac{d\tau}{2}\mathbf{k}_1, \tau_n + \frac{d\tau}{2}\right); \end{aligned} \tag{S41}$$

- fourth order Runge-Kutta (RK4)

$$
\begin{aligned}
\mathbf{x}_{n+1} &= \mathbf{x}_n + \frac{d\tau}{6}\left(\mathbf{k}_1 + 2\mathbf{k}_2 + 2\mathbf{k}_3 + \mathbf{k}_4\right), \\
\mathbf{k}_1 &= -\mathbf{v}(\tau_n, \mathbf{x}_n), \\
\mathbf{k}_2 &= -\mathbf{v}\left(\mathbf{x}_n + \frac{d\tau}{2}\mathbf{k}_1, \tau_n + \frac{d\tau}{2}\right), \\
\mathbf{k}_3 &= -\mathbf{v}\left(\mathbf{x}_n + \frac{d\tau}{2}\mathbf{k}_2, \tau_n + \frac{d\tau}{2}\right), \\
\mathbf{k}_4 &= -\mathbf{v}\left(\mathbf{x}_n + d\tau\mathbf{k}_3, \tau_n + d\tau\right).
\end{aligned}
\tag{S42}
$$

The time step $d\tau$ is adopted to integrate the ODE (S40) within the time interval $[0; t_{N_K} - t_1]$ and the superscript $n$ denotes the numerical solution at a given time step. $\tau \in [0; t_{N_K} - t_1]$ represents the local time coordinate which advances the solution in time. We choose $d\tau = 0.5$ which corresponds to approximately 3.5 times the time frame of acquisition (0.144 *sec*). This has been deliberately chosen in order to highlight even more the benefit of a high order reconstruction, because it contains much more information and thus can produce very good results even with large time steps which cover more than three consecutive frames, i.e. more than three adjacent computational cells. If classical linear tracking is used, i.e. $P1$ reconstruction is applied, the results suffer from lack of resolution (Fig. 2l) and the accuracy is drastically reduced.

#### S1.11 Particle directionality

For analyzing particle directionality, a direction vector $\mathbf{d} = (d_x, d_y, d_z)$ has to be defined. Specifically, this must be a unit length vector, so that it does not affect the projected velocity:

$$v_d = \mathbf{v} \cdot \mathbf{d}. \tag{S43}$$

Here, $v_d$ is the magnitude of the projected velocity along direction $\mathbf{d}$ and it is a scalar. $v_d$ has been used both for defining anterograde and retrograde as well as to get the information of the velocity component contributing to the movement in the direction $\mathbf{d}$ (i.e. directional velocity). Two different reference vectors are considered: i) $x-$direction (Option 1) and ii) axonal geometry (Option 2). The reference vector along the $x-$axis is simply given by $\mathbf{d} = (1, 0, 0)$. The axonal geometry is defined by a sequence of discrete points that are manually selected on top of the axon and the corresponding $(x, y, z)$ coordinates are retrieved by ImageJ tool. The reference line is then reconstructed by SHOT-R, from which one can easily extract the spatially dependent reference vector $\mathbf{d}(x, y, z)$ at any chosen point $\mathbf{x}_p = (x_p, y_p, z_p)$ by computing the derivative of the reference line, i.e. the tangent vector to the curve.

#### S1.12 Co-trafficking analysis

All particle trajectories for each channel undergo the SHOT-R algorithm, thus obtaining the corresponding high order reconstruction polynomials. Before carrying out the co-trafficking analysis, two preliminary filters are applied within the same axon: i) we exclude all pairs that do not share any temporal interval (by definition they cannot be co-trafficked); ii) we exclude all pairs which do not present at least 80% of the spatial path in common.

Once a pair passes the aforementioned thresholds, a common trajectory is then built. This is done by considering as input data all the spatiotemporal coordinates of each pair, so that the known information is maximized. The reconstruction polynomial is then computed according to a least square procedure without imposing any linear constraints, i.e. we directly solve the first overdetermined linear system in (S7) by means of classical normal equation technique. The unified trajectory constitutes the best solution which fits the known data by minimization of the square of the distances, thus it can be considered as the ideal co-trafficked path, if it exists. The errors are measured in $L_\infty$ norm for both particles of the pair and summed up along the entire common path in time. This is the reason why we set 1.2 $\mu m$ as $L_\infty$ threshold, which corresponds to the sum of the estimated diameters of particle pairs.

##### S1.13 Statistic analysis

All data are analyzed with Prism (GraphPad 6 or 7). For all tests, the significance level is $\alpha = 0.05$. Statistical significance is defined as: ns, not significant, * $P < 0.05$, ** $P < 0.01$, *** $P < 0.001$, **** $P < 0.0001$. The exact number of replicates and tests used are reported in Figure legends and further details on descriptive statistics are disclosed in Supplementary Table S3.

#### S2 Supplementary Figures and legends

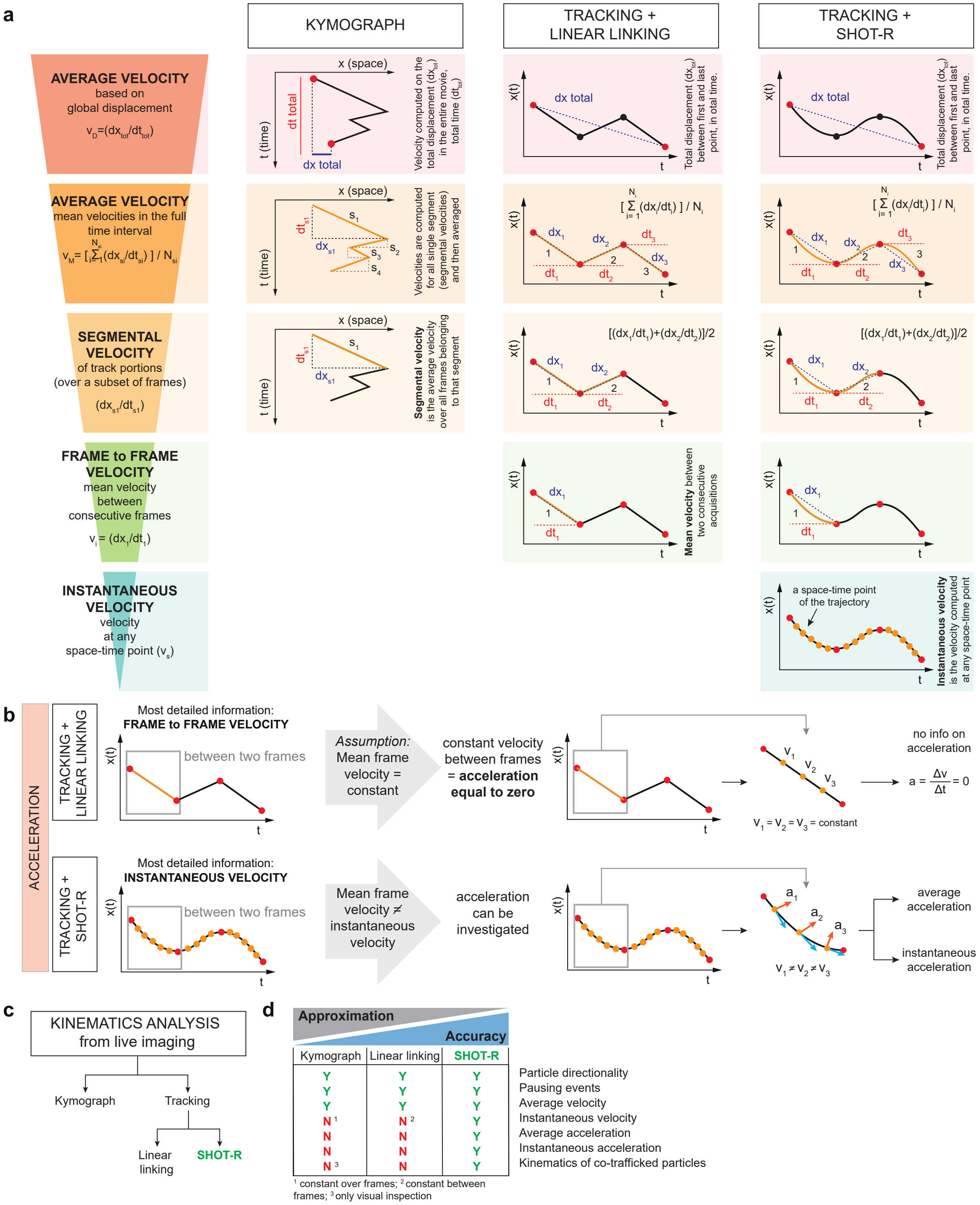


Figure S1: **Particle kinematics analysis: computing velocity and acceleration (a)** Schematic depicting how the existing methods to study particle kinematics (i.e. kymograph and single particle tracking

(SPT) combined with linear linking) and our new method (SHOT-R) compute velocity from average to more detailed information. The most detailed information achievable by each method corresponds to the last box of each column: segmental velocity for kymograph, frame to frame velocity for tracking and linear linking, and instantaneous velocity for tracking combined with SHOT-R. **(b)** Acceleration computed by tracking combined with linear linking (b upper panel) or with SHOT-R (b bottom panel). Linear linking assumes frame to frame constant velocity, meaning no acceleration (b upper panel); while curvilinear trajectory reconstruction by SHOT-R does not assume a constant frame to frame velocity and thus it allows to compute and investigate particle acceleration (b bottom panel). **(c-d)** Summary schematics of c) the three methods to study particles kinematic, d) the main advantages of SHOT-R over the existing methods: reduction of the approximation in particle trajectory reconstruction leading to a higher accuracy and the ability to study previously inaccessible information. Abbreviations: SHOT-R, Spatiotemporal High Order Trajectory Reconstruction.

---

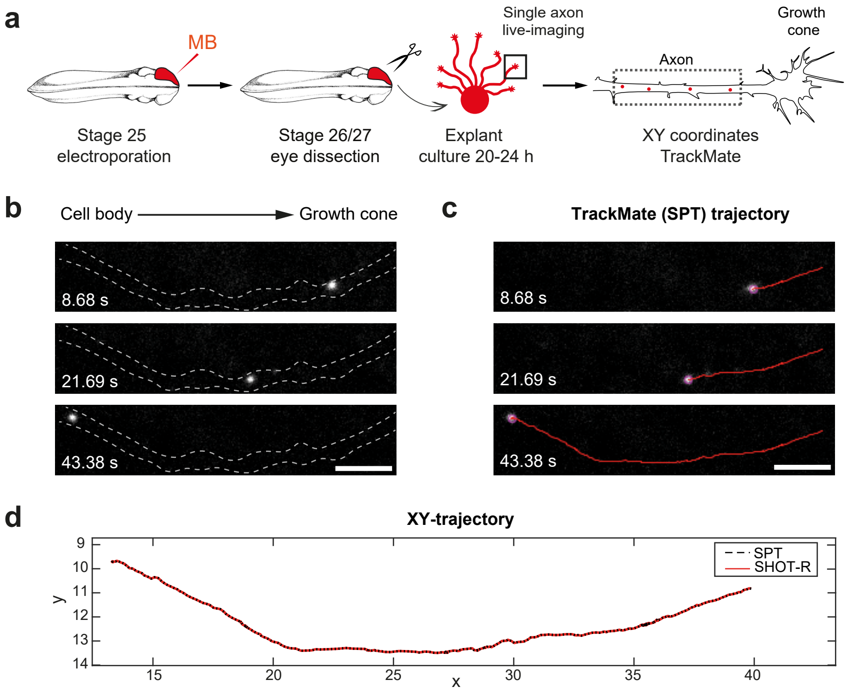


Figure S2: **SHOT-R initial condition: retrieve spatiotemporal particle coordinates (a)** Schematic of the experimental paradigm. **(b,c)** Representative time-lapse (b) and TrackMate trajectory (c) of a retrograde moving particle. The illustrative snapshots are 3 selected frames out of 300. **(d)** Comparison of SPT and SHOT-R in $xy$ trajectory reconstruction of the particle in panel b and c. Let observe that SPT and SHOT-R trajectories are overlapping. Abbreviations: MB, molecular beacon (recognizing the endogenous pre-miR-181a-1); SPT, single particle tracking; SHOT-R, Spatiotemporal High Order Trajectory Reconstruction. Scale bars: 5 $\mu m$ (b,c).

---

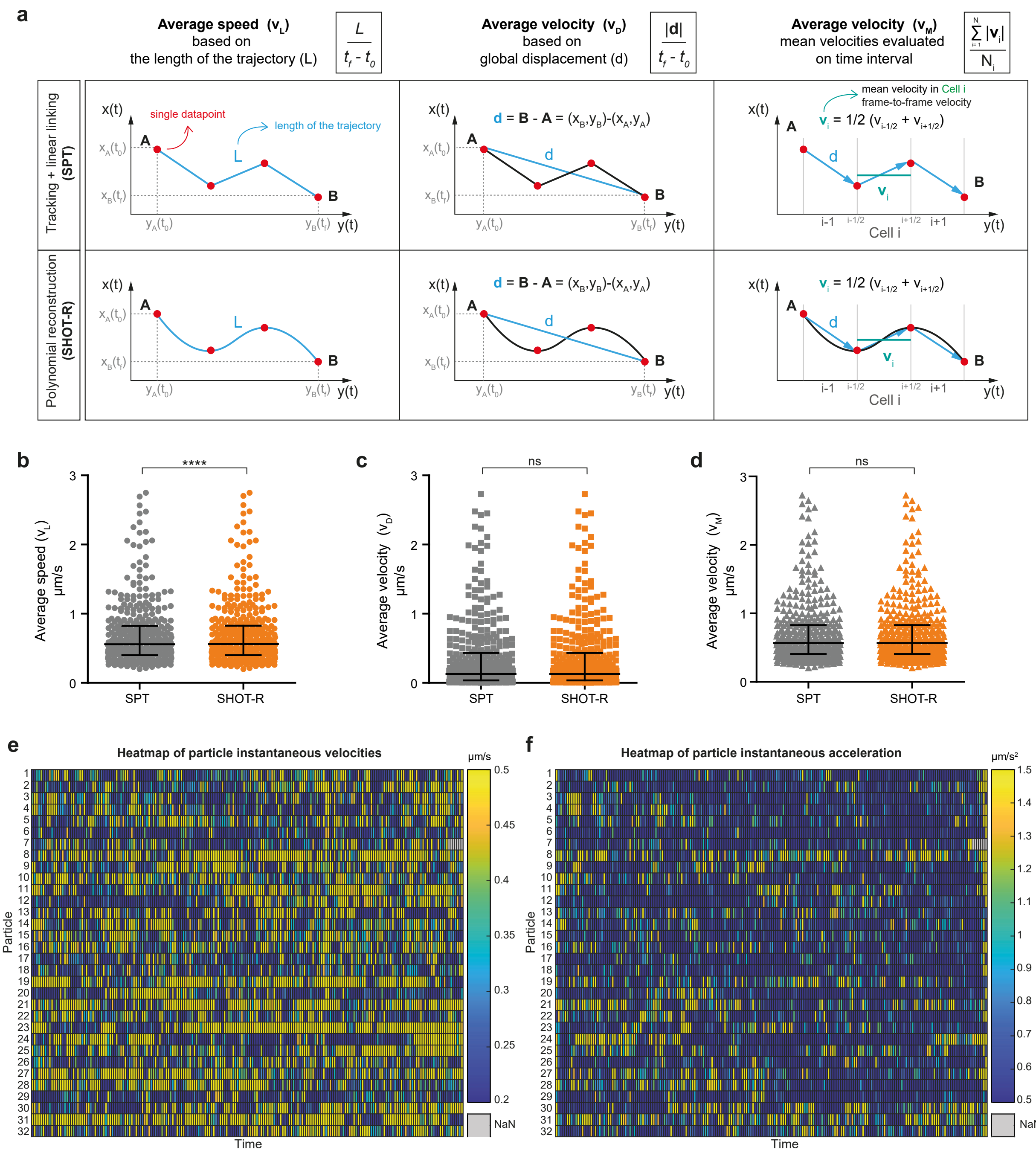


Figure S3: **SHOT-R: from average to instantaneous velocity and acceleration (a)** Schematics and formulae for average speed and velocity. Blue vectors are the local or global displacement used to compute velocity. **(b-d)** Velocities computed on linear (SPT) or polynomial (SHOT-R) reconstruction of the particle trajectory. (b) Average speed ($v_L$) based on the length trajectory. (c) Average velocity ($v_D$) based on particle global displacement. (d) Average velocity ($v_M$) based on the mean of all the $v_i$ for each time interval. **(e,f)** Heatmap of the instantaneous velocities (e) and acceleration (f) over time for the all particles with trajectories spanning at least 280 frames. Abbreviations: SPT, single particle tracking; SHOT-R, Spatiotemporal High Order Trajectory Reconstruction. Data information: ns, not significant, **** $P < 0.0001$. Values are median with interquartile range. Data are not normally distributed (Shapiro-Wilk normality test), two-tailed Wilcoxon matched pairs test (b-d). Total number of particles analyzed: 372 (b-d); total number of axons analysed: 21 (b-f).

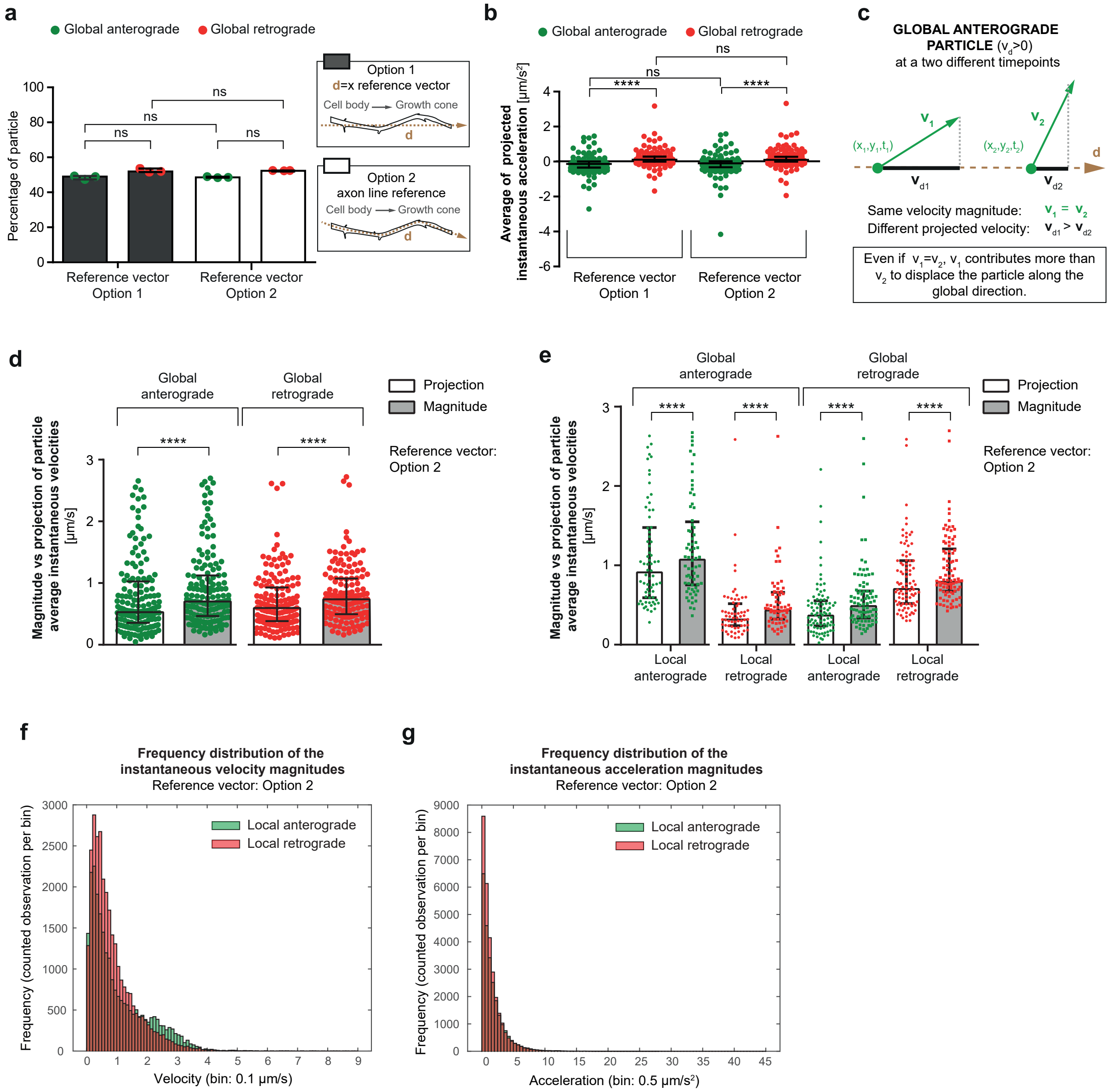


Figure S4: **SHOT-R: advances in reconstructing particle directionality (a)** Percentage of pre-miR-181a-1 anterograde and retrograde particles in respect to the reference **d**: $x$ vector (Option 1) or axon line (Option 2). **(b)** Average of directional instantaneous acceleration of global anterograde and retrograde particles in respect to the reference **d**: $x$ vector (Option 1) or axon line (Option 2). **(c)** Illustrative example of a particle moving at the same velocity ($v_1$ and $v_2$ magnitude) at two different time points ($t_1$ and $t_2$), but with a different directional velocity ($v_{d1}$ and $v_{d2}$). **(d,e)** Magnitude and directional average instantaneous velocities for global anterograde and retrograde particles (d) and for local anterograde and retrograde particles (e). **(f,g)** Frequency distribution of the instantaneous velocity (f) and of the instantaneous acceleration (g) magnitude. Data information: ns, not significant; **** $P < 0.0001$. Each data point corresponds to one independent experiment (a) or one particle (b,d,e). Values are median with interquartile range. Data are not normally distributed (Shapiro-Wilk normality test), Kruskal–Wallis test followed by Dunn's multiple comparison (a,b), two-tailed Wilcoxon matched-pairs test (d,e). Total number of puncta analysed: 158 (b,d,e) including 68 global anterograde particles and 90 global retrograde particles.

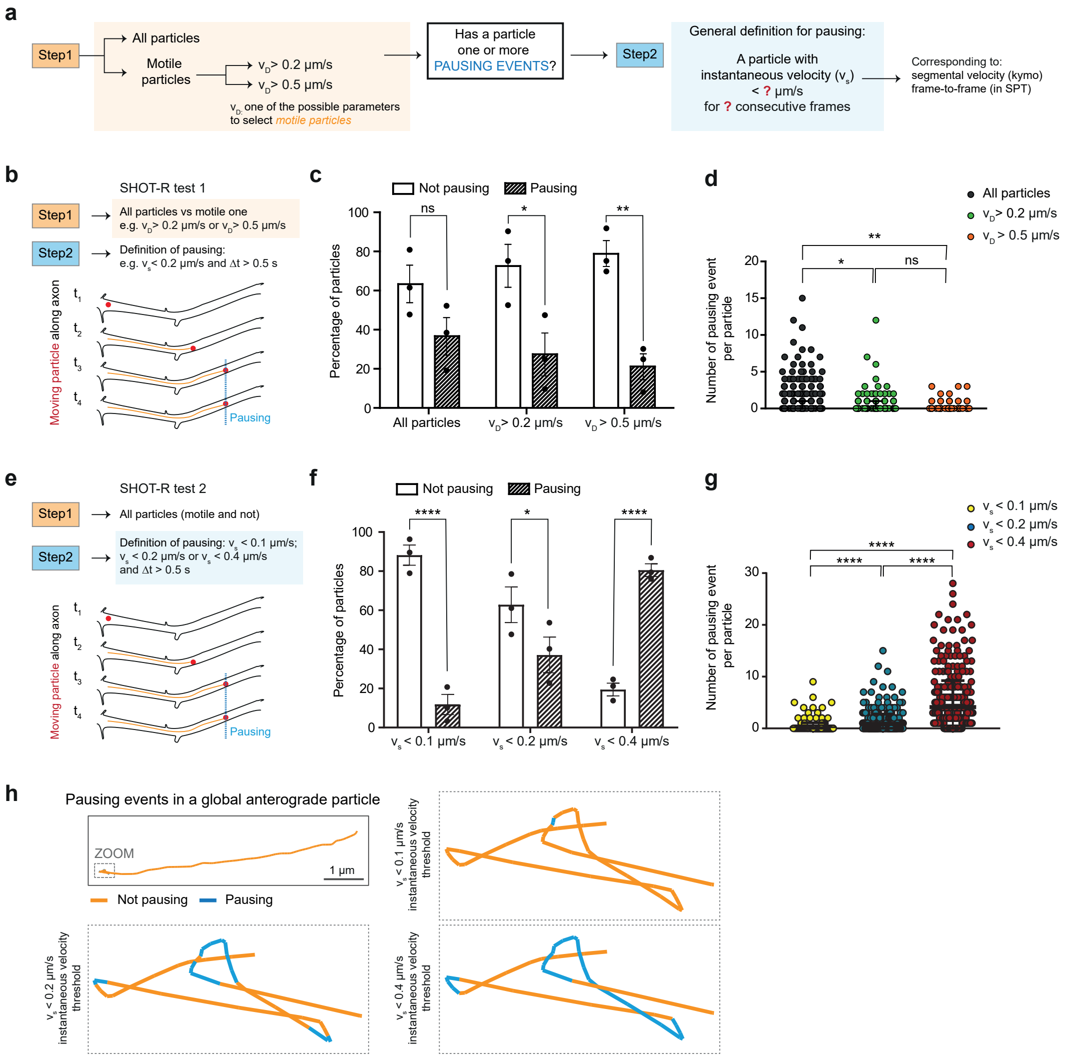


Figure S5: **SHOT-R: new advances in investigating pausing events (a)** Schematic of how pausing events are currently defined: selection or not of motile particle only (Step1) and threshold based on minimum frame number and on instantaneous velocity. **(b,e)** Experimental paradigm for SHOT-R analysis applying different parameters for Step1 and Step2. **(c,f)** Frequency distribution in percentage of pausing events per: (c) particle in the entire dataset (all particle) or in the subset of moving ($v_D > 0.2\ \mu m/s$) or fast moving particles ($v_D > 0.5\ \mu m/s$); (f) particle in the entire dataset (all particle) where different definitions of pauses were applied by thresholding the instantaneos velocity $v_s$. **(d,g)** Number of pausing events observed per: (d) particle trajectory in the entire dataset (all particle) or in subset of moving and fast moving particles; (g) particle in the entire dataset (all particle) where different definitions of pauses were applied by thresholding the instantaneos velocity $v_s$. **(h)** Representative trajectory (orange) interrupted by pauses (light blue): pauses change depending on the applied threshold for minimal instantaneos velocity $v_s$. Abbreviations: $v_D$, average velocity based on particle displacement; $v_s$, instantaneos velocity. Data information: ns, not significant; * P < 0.05; ** P < 0.01; **** P < 0.0001. Each data point corresponds to one particle (d,g), frequency in percentage was computed on n=3 independent experiment (c,f). Values are mean with SEM (c,f) median with interquartile range (d,g). Data are not normally distributed (Shapiro-Wilk normality test), 2-way ANOVA followed by Sidak's multiple comparison (c,f); Kruskal–Wallis test followed by Dunn's multiple comparison (d,g). Total number of particles analyzed: all particles (372, including 135 with pauses), moving particle $v_D > 0.2\ \mu m/s$ (158, including 42 with pauses) and fast moving particle $v_D > 0.5\ \mu m/s$ (85, including 17 with pauses). Total number of pausing events analyzed: all particles (341), moving particle $v_D > 0.2\ \mu m/s$ (83) and fast moving particle $v_D > 0.5\ \mu m/s$ (25).

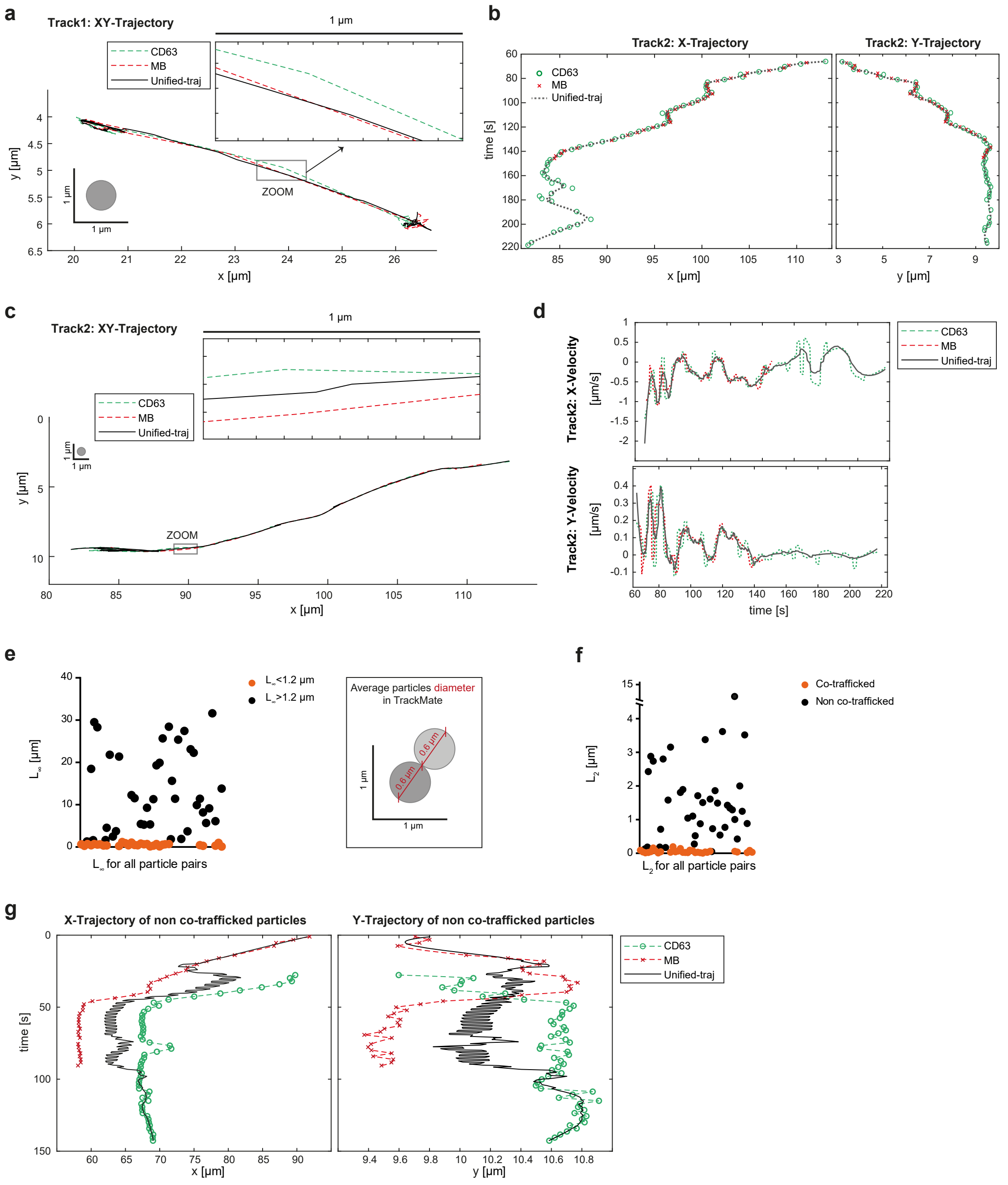


Figure S6: **SHOT-R: applications on quantitative co-trafficking analysis (a-c)** $xy$ trajectories of Track1 (a) and Track 2 (b,c) representative co-trafficked particles. The grey circle indicates a particle of 0.6 $\mu m$ in diameter as a reference. 1 $\mu m$ zoomed region highlights the accuracy of the unified trajectory over the single particle trajectories. **(d)** Representative instantaneous velocity plots for co-trafficked particles computed for Track 2. **(e)** $L_\infty$ norm for all particles pairs (CD63 and MB) analysed. $L_\infty$ threshold was set as 1.2 $\mu m$, corresponding to the sum of the estimated particle diameters in TrackMate (schematic in the right box). **(f)** $L_2$ norm for all particles pairs (CD63 and MB) analysed. **(g)** Representative $x$ and $y$ trajectory comparison against the unified reconstructed reference of non co-trafficked particles. Abbreviations: MB, molecular beacon (recognizing the endogenous pre-miR-181a-1, red); CD63, tetraspanin marker of late endosome/lysosome (green); Unified-traj, unified reference trajectory. Data information: Each data point corresponds to a CD63-MB pair-comparison. 3 independent experiments, 9 axons. Total number of particles analyzed: MB-pre-miR-181a-1 (87), CD63 (69), co-trafficked particle-pairs (31).

### S3 Supplementary Tables

Table S1: Numerical convergence rates obtained with high order accurate reconstruction of the velocity field given by (S36). Errors in $L_1$, $L_2$ and $L_\infty$ norms are reported for all position components $(x, y, z)$ with associated order of accuracy $\mathcal{O}$ on a sequence of refined mesh with characteristic size $\Delta t$.

**N = 1**

| | horizontal position $x$ | | | | | | horizontal position $y$ | | | | | | horizontal position $z$ | | | | | |
|---|---|---|---|---|---|---|---|---|---|---|---|---|---|---|---|---|---|---|
| $\Delta t$ | $\varepsilon_{L_1}$ | $\mathcal{O}(L_1)$ | $\varepsilon_{L_2}$ | $\mathcal{O}(L_2)$ | $\varepsilon_{L_\infty}$ | $\mathcal{O}(L_\infty)$ | $\varepsilon_{L_1}$ | $\mathcal{O}(L_1)$ | $\varepsilon_{L_2}$ | $\mathcal{O}(L_2)$ | $\varepsilon_{L_\infty}$ | $\mathcal{O}(L_\infty)$ | $\varepsilon_{L_1}$ | $\mathcal{O}(L_1)$ | $\varepsilon_{L_2}$ | $\mathcal{O}(L_2)$ | $\varepsilon_{L_\infty}$ | $\mathcal{O}(L_\infty)$ |
| 1.00e-2 | 1.89e-3 | - | 1.49e-3 | - | 1.64e-3 | - | 5.06e-3 | - | 4.00e-3 | - | 4.60e-3 | - | 7.74e-3 | - | 6.24e-3 | - | 7.62e-3 | - |
| 5.00e-3 | 4.73e-4 | 2.00 | 3.72e-4 | 2.00 | 4.11e-4 | 2.00 | 1.27e-3 | 2.00 | 1.00e-3 | 2.00 | 1.15e-3 | 2.00 | 1.94e-3 | 2.00 | 1.56e-3 | 2.00 | 1.91e-3 | 2.00 |
| 2.50e-3 | 1.18e-4 | 2.00 | 9.31e-5 | 2.00 | 1.03e-4 | 2.00 | 3.16e-4 | 2.00 | 2.50e-4 | 2.00 | 2.88e-4 | 2.00 | 4.84e-4 | 2.00 | 3.90e-4 | 2.00 | 4.77e-4 | 2.00 |
| 1.25e-3 | 2.95e-5 | 2.00 | 2.33e-5 | 2.00 | 2.57e-5 | 2.00 | 7.91e-5 | 2.00 | 6.25e-5 | 2.00 | 7.20e-5 | 2.00 | 1.21e-4 | 2.00 | 9.75e-5 | 2.00 | 1.19e-4 | 2.00 |

**N = 2**

| | horizontal position $x$ | | | | | | horizontal position $y$ | | | | | | horizontal position $z$ | | | | | |
|---|---|---|---|---|---|---|---|---|---|---|---|---|---|---|---|---|---|---|
| $\Delta t$ | $\varepsilon_{L_1}$ | $\mathcal{O}(L_1)$ | $\varepsilon_{L_2}$ | $\mathcal{O}(L_2)$ | $\varepsilon_{L_\infty}$ | $\mathcal{O}(L_\infty)$ | $\varepsilon_{L_1}$ | $\mathcal{O}(L_1)$ | $\varepsilon_{L_2}$ | $\mathcal{O}(L_2)$ | $\varepsilon_{L_\infty}$ | $\mathcal{O}(L_\infty)$ | $\varepsilon_{L_1}$ | $\mathcal{O}(L_1)$ | $\varepsilon_{L_2}$ | $\mathcal{O}(L_2)$ | $\varepsilon_{L_\infty}$ | $\mathcal{O}(L_\infty)$ |
| 1.00e-2 | 2.80e-4 | - | 2.49e-4 | - | 6.16e-4 | - | 4.86e-4 | - | 4.14e-4 | - | 5.91e-4 | - | 1.11e-3 | - | 9.49e-4 | - | 1.38e-3 | - |
| 5.00e-3 | 3.43e-5 | 3.03 | 3.02e-5 | 3.04 | 8.16e-5 | 2.92 | 5.96e-5 | 3.03 | 5.13e-5 | 3.01 | 7.39e-5 | 3.00 | 1.35e-4 | 3.04 | 1.16e-4 | 3.03 | 1.72e-4 | 3.00 |
| 2.50e-3 | 4.23e-6 | 3.02 | 3.69e-6 | 3.03 | 1.03e-5 | 2.98 | 7.42e-6 | 3.01 | 6.41e-6 | 3.00 | 9.24e-6 | 3.00 | 1.68e-5 | 3.01 | 1.45e-5 | 3.01 | 2.15e-5 | 3.00 |
| 1.25e-3 | 5.25e-7 | 3.01 | 4.55e-7 | 3.02 | 1.30e-6 | 2.99 | 9.27e-7 | 3.00 | 8.01e-7 | 3.00 | 1.15e-6 | 3.00 | 2.09e-6 | 3.00 | 1.81e-6 | 3.00 | 2.69e-6 | 3.00 |

**N = 3**

| | horizontal position $x$ | | | | | | horizontal position $y$ | | | | | | horizontal position $z$ | | | | | |
|---|---|---|---|---|---|---|---|---|---|---|---|---|---|---|---|---|---|---|
| $\Delta t$ | $\varepsilon_{L_1}$ | $\mathcal{O}(L_1)$ | $\varepsilon_{L_2}$ | $\mathcal{O}(L_2)$ | $\varepsilon_{L_\infty}$ | $\mathcal{O}(L_\infty)$ | $\varepsilon_{L_1}$ | $\mathcal{O}(L_1)$ | $\varepsilon_{L_2}$ | $\mathcal{O}(L_2)$ | $\varepsilon_{L_\infty}$ | $\mathcal{O}(L_\infty)$ | $\varepsilon_{L_1}$ | $\mathcal{O}(L_1)$ | $\varepsilon_{L_2}$ | $\mathcal{O}(L_2)$ | $\varepsilon_{L_\infty}$ | $\mathcal{O}(L_\infty)$ |
| 1.00e-2 | 7.66e-5 | - | 6.60e-5 | - | 1.04e-4 | - | 9.05e-5 | - | 8.24e-5 | - | 2.27e-4 | - | 2.99e-4 | - | 2.69e-4 | - | 6.88e-4 | - |
| 5.00e-3 | 4.68e-6 | 4.03 | 4.01e-6 | 4.04 | 4.94e-6 | 4.39 | 5.61e-6 | 4.01 | 5.00e-6 | 4.04 | 1.50e-5 | 3.92 | 1.88e-5 | 3.99 | 1.68e-5 | 4.00 | 4.94e-5 | 3.80 |
| 2.50e-3 | 2.90e-7 | 4.01 | 2.50e-7 | 4.00 | 3.10e-7 | 3.99 | 3.47e-7 | 4.01 | 3.05e-7 | 4.04 | 9.51e-7 | 3.98 | 1.17e-6 | 4.01 | 1.03e-6 | 4.03 | 3.19e-6 | 3.95 |
| 1.25e-3 | 1.81e-8 | 4.00 | 1.56e-8 | 4.00 | 1.94e-8 | 4.00 | 2.16e-8 | 4.01 | 1.88e-8 | 4.02 | 5.96e-8 | 4.00 | 7.29e-8 | 4.01 | 6.34e-8 | 4.02 | 2.01e-7 | 3.99 |

**N = 4**

| | horizontal position $x$ | | | | | | horizontal position $y$ | | | | | | horizontal position $z$ | | | | | |
|---|---|---|---|---|---|---|---|---|---|---|---|---|---|---|---|---|---|---|
| $\Delta t$ | $\varepsilon_{L_1}$ | $\mathcal{O}(L_1)$ | $\varepsilon_{L_2}$ | $\mathcal{O}(L_2)$ | $\varepsilon_{L_\infty}$ | $\mathcal{O}(L_\infty)$ | $\varepsilon_{L_1}$ | $\mathcal{O}(L_1)$ | $\varepsilon_{L_2}$ | $\mathcal{O}(L_2)$ | $\varepsilon_{L_\infty}$ | $\mathcal{O}(L_\infty)$ | $\varepsilon_{L_1}$ | $\mathcal{O}(L_1)$ | $\varepsilon_{L_2}$ | $\mathcal{O}(L_2)$ | $\varepsilon_{L_\infty}$ | $\mathcal{O}(L_\infty)$ |
| 1.00e-2 | 1.18e-5 | - | 1.17e-5 | - | 4.77e-5 | - | 9.72e-6 | - | 8.65e-6 | - | 2.74e-5 | - | 5.11e-5 | - | 4.89e-5 | - | 1.88e-4 | - |
| 5.00e-3 | 3.68e-7 | 5.01 | 3.67e-7 | 4.99 | 1.98e-6 | 4.59 | 2.76e-7 | 5.14 | 2.36e-7 | 5.20 | 4.66e-7 | 5.88 | 1.42e-6 | 5.17 | 1.23e-6 | 5.32 | 3.31e-6 | 5.83 |
| 2.50e-3 | 1.11e-8 | 5.05 | 1.06e-8 | 5.12 | 6.60e-8 | 4.91 | 8.40e-9 | 5.04 | 7.21e-9 | 5.03 | 9.98e-9 | 5.55 | 4.27e-8 | 5.06 | 3.66e-8 | 5.07 | 5.48e-8 | 5.92 |
| 1.25e-3 | 3.38e-10 | 5.04 | 3.09e-10 | 5.10 | 2.09e-9 | 4.98 | 2.61e-10 | 5.01 | 2.25e-10 | 5.00 | 3.12e-10 | 5.00 | 1.32e-9 | 5.01 | 1.14e-9 | 5.01 | 1.57e-9 | 5.12 |

**N = 5**

| | horizontal position $x$ | | | | | | horizontal position $y$ | | | | | | horizontal position $z$ | | | | | |
|---|---|---|---|---|---|---|---|---|---|---|---|---|---|---|---|---|---|---|
| $\Delta t$ | $\varepsilon_{L_1}$ | $\mathcal{O}(L_1)$ | $\varepsilon_{L_2}$ | $\mathcal{O}(L_2)$ | $\varepsilon_{L_\infty}$ | $\mathcal{O}(L_\infty)$ | $\varepsilon_{L_1}$ | $\mathcal{O}(L_1)$ | $\varepsilon_{L_2}$ | $\mathcal{O}(L_2)$ | $\varepsilon_{L_\infty}$ | $\mathcal{O}(L_\infty)$ | $\varepsilon_{L_1}$ | $\mathcal{O}(L_1)$ | $\varepsilon_{L_2}$ | $\mathcal{O}(L_2)$ | $\varepsilon_{L_\infty}$ | $\mathcal{O}(L_\infty)$ |
| 1.00e-2 | 3.90e-6 | - | 3.91e-6 | - | 1.71e-5 | - | 1.92e-6 | - | 1.98e-6 | - | 9.03e-6 | - | 1.35e-5 | - | 1.29e-5 | - | 4.73e-5 | - |
| 5.00e-3 | 5.63e-8 | 6.12 | 4.93e-8 | 6.31 | 1.57e-7 | 6.77 | 2.99e-8 | 6.01 | 2.99e-8 | 6.05 | 1.70e-7 | 5.73 | 2.24e-7 | 5.91 | 2.20e-7 | 5.87 | 1.19e-6 | 5.31 |
| 2.50e-3 | 8.54e-10 | 6.04 | 7.33e-10 | 6.07 | 1.27e-9 | 6.94 | 4.58e-10 | 6.03 | 4.32e-10 | 6.11 | 2.77e-9 | 5.94 | 3.47e-9 | 6.02 | 3.26e-9 | 6.07 | 2.07e-8 | 5.85 |
| 1.25e-3 | 1.32e-11 | 6.01 | 1.14e-11 | 6.01 | 1.49e-11 | 6.42 | 7.05e-12 | 6.02 | 6.39e-12 | 6.08 | 4.39e-11 | 5.98 | 5.36e-11 | 6.02 | 4.85e-11 | 6.07 | 3.31e-10 | 5.97 |

Table S2: Mathematical test with initial condition (S37)-(S38). Errors with respect to the analytical solution are measured in $L_1$, $L_2$ and $L_\infty$ norms for $(x, y)$ trajectory and velocity. Second order ($P1$, SPT) and fourth order ($P3$, SHOT-R) accurate reconstructions are used, demonstrating that the high order results systematically exhibit a lower error. The errors listed in Table S2 are used to construct the plots depicted in Fig. 1g.

| **Norm** | **Mesh** | $x$ $P1$ | $x$ $P3$ | $y$ $P1$ | $y$ $P3$ | $v_x$ $P1$ | $v_x$ $P3$ | $v_y$ $P1$ | $v_y$ $P3$ |
|---|---|---|---|---|---|---|---|---|---|
| $L_1$ | $\Delta t_1$ | 7.41e-4 | 1.17e-4 | 1.85e-2 | 4.17e-3 | 3.47e-2 | 5.50e-3 | 8.72e-1 | 1.97e-1 |
| | $\Delta t_2$ | 1.85e-4 | 8.34e-6 | 4.66e-3 | 2.42e-4 | 1.74e-2 | 7.86e-4 | 4.37e-1 | 2.29e-2 |
| | $\Delta t_3$ | 4.63e-5 | 5.41e-7 | 1.16e-3 | 1.44e-5 | 8.69e-3 | 1.02e-4 | 2.18e-1 | 2.72e-3 |
| $L_2$ | $\Delta t_1$ | 1.09e-6 | 2.27e-8 | 2.55e-4 | 1.54e-5 | 2.46e-3 | 5.12e-5 | 5.74e-1 | 3.50e-2 |
| | $\Delta t_2$ | 6.85e-8 | 1.28e-10 | 1.60e-5 | 4.39e-8 | 6.17e-4 | 1.15e-6 | 1.44e-1 | 3.96e-4 |
| | $\Delta t_3$ | 4.29e-9 | 5.59E-13 | 1.00e-6 | 1.51e-10 | 1.54e-4 | 2.01e-8 | 3.61e-2 | 5.43e-6 |
| $L_\infty$ | $\Delta t_1$ | 2.39e-3 | 4.16e-4 | 1.93e-2 | 8.05e-3 | 1.41e-1 | 2.47e-2 | 1.13e-0 | 4.96e-1 |
| | $\Delta t_2$ | 6.10e-4 | 3.42e-5 | 4.84e-3 | 3.45e-4 | 7.14e-2 | 4.04e-3 | 5.66e-1 | 4.24e-2 |
| | $\Delta t_3$ | 1.53e-4 | 2.33e-6 | 1.21e-3 | 1.49e-5 | 3.58e-2 | 5.48e-4 | 2.83e-1 | 3.50e-3 |
